# Foliar fungal symbionts in sympatric yellow monkeyflowers along elevation gradients in the Sierra Nevada

**DOI:** 10.64898/2026.01.18.700187

**Authors:** Bolívar Aponte Rolón, Kathleen G. Ferris, Sunshine A. Van Bael

## Abstract

Microbial symbionts have the potential to contribute to host-plant’s ecological and evolutionary success, especially in plants’ adaptions to harsh environments, however their role has often been overlooked. Conversely, how host local adaptation (e.g., trait divergence across elevation) shapes the composition of associated microbial symbiont communities remains poorly understood. We explored how foliar endophytic fungi (FEF) abundance, richness, and community composition in three sympatric Monkeyflowers, an ecologically diverse group of flowering plants, change across elevation in the Sierra Nevada, CA, USA. We asked: Q1) Are there differences in leaf functional traits and FEF communities among sympatric *Mimulus* species populations at similar elevations? Q2) How do traits and FEF communities change across elevation within species? Q3) Are FEF richness, diversity and community composition correlated with leaf functional traits and/or elevation? Q4) How does FEF community composition differ with geographic distance within each species? We collected *M. guttatus*, *M. nasutus*, and *M.laciniatus* individuals from natural populations across the Sierra Nevada, measured leaf functional traits and used ITS sequencing to describe the leaf endophyte community. We found significant associations of FEF community composition with host species, and elevation, suggesting that these factors influence fungal community composition. Furthermore, FEF community dissimilarity was correlated with geographic distance indicating isolation by distance and limited dispersal of fungal endophytes. We detected the prevalence of *Vishniacozyma victoriae*, an endophyte found most commonly in Antarctica, across all Sierran *Mimulus* populations. The presence of *V. victoriae* could play a role in the adaptation of *Mimulus* to cold, high elevation environments. Our findings offer novel insights into the intricate interactions between fungal endophyte communities, plant traits, and elevation.

## 2. Introduction

Microbial dispersal is challenging to trace. The general assumption is that most, if not all, microbes are ubiquitous. Nevertheless, the dispersal of microbial communities is not random as it is the result of neutral and selective processes influenced by environmental conditions, geographic distance, and host species (Rebolleda Gómez and Ashman, 2019). The co-evolution and symbiotic interactions plant microbes, such as fungi (Remy et al., 1994; Wang and Qiu, 2006; Field et al., 2015; reviewed in Peay et al., 2016) and bacteria (Soltis et al., 1995; Adams, 2002; Delaux et al., 2015), aided plants’ colonization of land across multiple varying climates. However, little is still known about how plant microbiomes influence or are influenced by local adaptation of their host species. Geographic location and environmental conditions are known to influence community composition of symbiotic fungi, with varying effects across different fungal guilds (Bowman and Arnold, 2021; Kivlin et al., 2022). Previous studies have suggested that foliar endophytic fungi (FEF), which live inside plant leaves, can alter plant traits under stressful conditions such as drought (Giauque and Hawkes, 2013; Song et al., 2016; Giauque et al., 2019) and therefore may contribute to local adaptation. Given these close associations between host and symbiont the resulting FEF community is influenced by the interplay of environmental conditions, geographic distance and host species. Host traits and host identity can influence FEF communities in a variety of ways. Leaf functional traits, such as leaf thickness, leaf mass per area, and leaf toughness are known to influence plant’s response to abiotic factors like light, temperature, and hydraulic constraints (Nicotra et al., 2011; Oguchi et al., 2018). These traits serve as host-imposed filters that play a role in structuring FEF communities (Saunders et al., 2010; Tellez et al., 2022). Jointly, host identity has been shown to structure the foliar microbiomes of plants. For example, a study of the classical plant model, *Arabidopsis thaliana*, revealed that bacterial and fungal communities in leaves were shaped by host genotype (Horton et al., 2014). Recently, Karasov et al. (2024), found robust trends in further support of this hypothesis, showing that the bacterial leaf microbiome in diverse European populations of *A. thaliana* is influenced by genetic variation in the host. At the local scale, *A. thaliana* genotype can impose strong selection on the composition of its leaf microbiome, but at the continental scale, while genotype still plays a role, geography and abiotic factors also contribute strongly to symbiont community composition (Karasov et al., 2024). The patterns observed in this model system encourage testing these hypotheses in other plant species to determine their broader applicability. The *Mimulus guttatus* species complex, a closely related interfertile group of species with considerable morphological and ecological variation, is an ideal model system to explore questions about FEF communities and their relationship with host species traits and environmental conditions (Vickery, 1978; Wu et al., 2008).

Previous studies have examined floral and root microbial symbionts in *Mimulus* species, yet little is known about their foliar symbionts (Belisle et al., 2012; Rebolleda Gómez and Ashman, 2019; McIntosh et al., 2024). Aboveground, Beslile et al. (2012) reported on the distribution of diverse fungal communities in the flower nectar of *Mimulus aurantiacus*. The authors considered flowers as islands in a metapopulation system and found that the frequency of micro-fungi (i.e., yeasts) was significantly correlated to the location of the plant and marginally correlated with the density of the flowers in the plant (Belisle et al., 2012). Another study by Rebolleda et al (2019), examined epiphytic bacterial communities of different flower parts in *Mimulus guttatus* and found that pollinator choice drives strong micro-environmental selection of bacterial communities, overwhelming the effects of geographic distance. Belowground, arbuscular mycorrhizal fungi (AMF) are known to associate with *M. guttatus*.

Foliar symbionts can increase plant fitness under stressful conditions (Giauque et al., 2019; Shankar Naik, 2019; Aimone et al., 2023; Karasov et al., 2024). Altitudinal gradients provide excellent natural laboratories to explore environmental effects on plant-microbe interactions. The *M. guttatus* species complex occupies a wide range of habitats in Western North America (Wu et al., 2008), including elevational gradients which makes it an interesting system to examine the composition of FEF communities across species and determine if they align with broader ecological patterns. A study by Kazenel et al. (2019), found that the composition of root and leaf endophytes in cool-season, perennial grasses, varied differently with altitude and warming. Others have found that elevation and site are the strongest predictors of fungal taxa (Cordier et al., 2012). To our knowledge, no previous studies have considered symbionts in leaf tissue of *Mimulus* or how sympatric host species and elevational gradients influence symbiont community composition in the *Mimulus guttatus* species complex.

In this study we use amplicon sequencing to explore how FEF communities vary among sympatric populations of *M. guttatus*, *M. laciniatus* and *M. nasutus* collected along replicated elevation gradients throughout the central Sierra Nevada, CA (Fig. 1). We test the following questions: Q1) Are there differences in leaf functional traits among sympatric *Mimulus* species populations at similar elevations? Q2) Within each host species, how do leaf traits vary along the elevation gradient? Q3) To what extent do leaf traits and elevation explain variation in FEF richness, diversity, composition, and community structure across individuals and species? Q4) How does FEF community composition differ with geographic distance within each species? We include leaf thickness (LT), leaf punch strength (LPS), leaf mass per area (LMA), and anthocyanin content index (ACI) because these traits are known to influence the structure of FEF communities and play key roles in plant defenses against herbivores and pathogens (Gould, 2004; Saunders et al., 2010; Tellez et al., 2022; Aponte Rolón et al., 2024). Additionally, we include leaf lobe index (LBI) considering how distinct leaf shape is between the three *Mimulus* species and due to its potential effects on light exposure, temperature regulation, and hydraulic constraints within leaf tissue, which may, in turn, impact FEF symbionts (Nobel, 2009; Ferris et al., 2015). We predicted that abundance, diversity,and richness would decline with increased elevation, and community composition of FEF would be more similar among the same sites (alpha diversity) than between sites (beta diversity) regardless of host species. With these questions we tested whether FEF community responses to altitude were consistent across host species. Our results provide insights into how host genotype and environmental variation affect the ecological dynamics of plant-microbe interactions.

**Figure 1:**
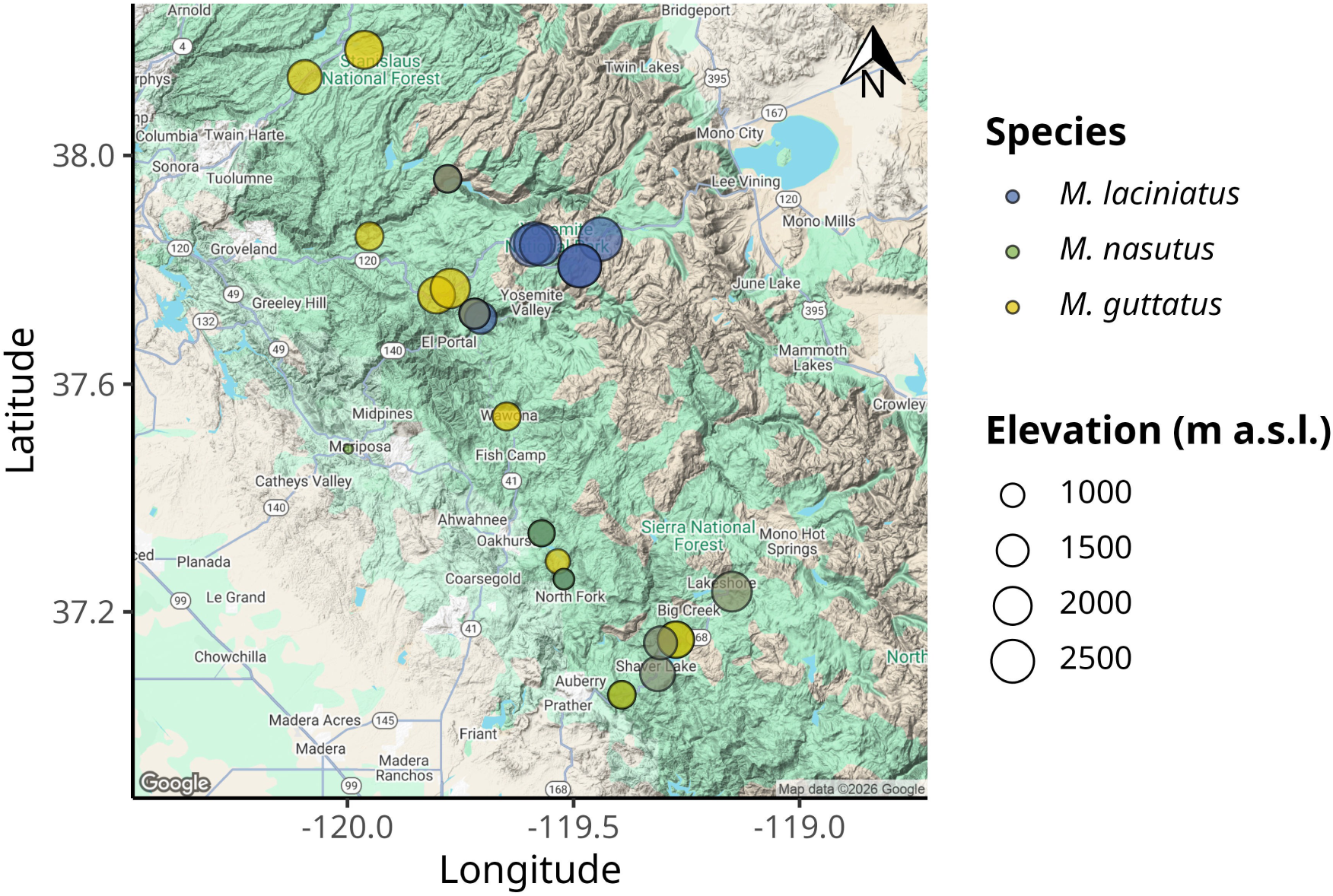
Map of *Mimulus* spp. sampling sites in the Sierra Nevada, California, USA. A) Map of the Sierra Nevada region showing the location of the three species sampled. The color gradient represents the elevation gradient from low (yellow) to high (purple).

## 3. Materials and Methods

### 3.1 Sample Collection

To examine the effects of geography and host species on FEF communities, we collected plant specimens during April-July 2021 and 2022 from populations of *M. guttatus*, *M. laciniatus*, and *M. nasutus* (syn. *Erythranthe guttata*, *Erythranthe laciniata*, and *Erythranthe nasuta*) across Stanislaus National Forest (SNF), Sierra National Forest (SINF) and Yosemite National Park (YNP), CA, USA. We haphazardly selected sites close to the main roads based on the presence of a viable population with at least ∼ 50 individuals per species. Samples collected from YNP were collected from non-wilderness areas on the side of the road. We determined population viability ensuring that they had individuals flowering or close to flowering stage. We collected between 5 - 20 individuals per species from a total of 25 sites (Table 1). We selected individuals that possessed healthy looking leaves and no visible signs of pathogen damage or senescence. At sites where two species were present, we collected individuals that were at least ∼ 25 meters apart. We collected sample specimens by carefully uprooting the plant and placing into individual plastic bags (e.g., Ziploc®) and preserving in an ice chest until return to the field laboratory at the UC Merced Yosemite Field Station, YNP, CA, USA. Plant specimens were processed within 8 hrs of collection.

**Table 1.**

|  | Model 1 |  | Model 2 |  | Model 3 |  | Model 4 |  |
| --- | --- | --- | --- | --- | --- | --- | --- | --- |
| Predictors | Estimate | CI [t-statistic] | Estimate | CI [t-statistic] | Estimate | CI [t-statistic] | Estimate | CI [t-statistic] |
| Intercept | 0.59 *** | (0.49 - 0.69 )<br>[11.35] | 0.83 *** | (0.75 - 0.91 )<br>[20.11] | 0.78 *** | (0.74 - 0.82 )<br>[38.06] | 0.86 *** | (0.83 - 0.89 )<br>[48.65] |
| *M. nasutus* | -0.01 ** | (-0.02 - -0.00 )<br>[-2.93] | 0.02 *** | (0.01 - 0.02 )<br>[3.72] | 0.02 *** | (0.01 - 0.03 )<br>[3.91] |  |  |
| *M. guttatus* | 0.02 *** | (0.01 - 0.03 )<br>[4.19] | 0.04 *** | (0.03 - 0.05 )<br>[8.74] | 0.04 *** | (0.03 - 0.05 )<br>[8.93] |  |  |
| Elevation | -0.00 | (-0.00 - 0.00 )<br>[-1.64] | -0.00 | (-0.00 - 0.00 )<br>[-1.36] |  |  |  |  |
| ACI | 0.05 *** | (0.03 - 0.06 )<br>[7.03] | -0.08 *** | (-0.08 - -0.07 )<br>[-29.93] | -0.08 *** | (-0.08 - -0.07 )<br>[-29.90] | -0.08 *** | (-0.08 - -0.07 )<br>[-32.48] |
| LT | 0.03 * | (0.00 - 0.06 )<br>[2.14] | -0.03 *** | (-0.04 - -0.02 )<br>[-5.68] | -0.03 *** | (-0.04 - -0.02 )<br>[-5.71] | -0.05 *** | (-0.06 - -0.04 )<br>[-9.51] |
| LPS | -0.03 *** | (-0.05 - -0.01 )<br>[-3.42] | -0.02 ** | (-0.03 - -0.00 )<br>[-2.65] | -0.02 ** | (-0.03 - -0.00 )<br>[-2.64] |  |  |
| LMA | -0.04 *** | (-0.06 - -0.03 )<br>[-5.75] | 0.02 *** | (0.01 - 0.03 )<br>[4.83] | 0.02 *** | (0.01 - 0.03 )<br>[4.88] | 0.01 *** | (0.01 - 0.02 )<br>[3.74] |
| LBI | -0.01 * | (-0.02 - -0.00 )<br>[-2.53] | -0.01 * | (-0.02 - -0.00 )<br>[-2.34] | -0.01 * | (-0.02 - -0.00 )<br>[-2.41] | -0.00 | (-0.01 - 0.01 )<br>[-0.22] |
| *Random effects* |  |  |  |  |  |  |  |  |
| N | 4200 |  | 4200 |  | 4200 |  | 4200 |  |
| logLik | 6729.85 |  | 8154.59 |  | 8163.50 |  | 8330.81 |  |
| AIC | -13437.71 |  | -16249.18 |  | -16269.01 |  | -16603.63 |  |

Significance levels are represented by asterisks [p < 0.05 (\*), p < 0.01 (\*\*), p < 0.001 (\*\*\*), and p < 0.0001 (\*\*\*\*)]. T statistics in brackets.

### 3.2 Leaf trait measurements

To quantify phenotypic diversity among and within these species, we measured the following leaf traits: leaf thickness (LT), leaf punch strength (LPS), leaf mass per area (LMA), anthocyanin content index (ACI) and leaf lobe index (LBI). We cleaned plants with tap water to remove all soil and debris remnants from the leaves and roots. We removed all healthy leaves (∼ 5 - 10) from the stems and took three measurements per trait from three haphazardly selected leaves from individual plants, except for LBI, where we measured only one leaf per plant. To calculate LBI we used a transparency film to hold the leaf in place and flatten, after which we took a digital photograph and then followed Ferris et al. (2015). Briefly, leaf lobing is calculated as the convex hull area minus the true leaf area divided by convex hull area of leaf in ImageJ [v1.52r; Schneider et al. (2012)]. We measured ACI content with ACM-200plus (Opti-Sciences Inc. Hudson, New Hampshire, U.S.A.) on haphazardly selected locations of the leaf surface (working from the petiole out to the leaf tip) (Tellez et al., 2022). The ACM-200 calculates an ACI value from the ratio of % transmittance at 931 nm/% transmittance at 525 nm (n.d.), effectively accounting for leaf thickness. We measured LT (μm) with a Mitutoyo 7327 Micrometer Gauge (Mitutoyo, Takatsu-ku, Kawasaki, Japan) on haphazardly selected locations of the leaf lamina, taking care to avoid major and secondary veins. We used an Imada DST-11a digital force gauge (Imada Inc., Northbrook, IL, United States) to measure LPS, a measure of leaf toughness, on the lamina of each leaf selected, avoiding minor leaf veins when possible (Tellez et al., 2022). It functions by conducting punch-and-die tests with a sharp-edged cylindrical steel punch (2.0 mm diameter) and a steel die with a sharp-edged aperture of small clearance (0.05 mm). Once LPS was measured, we used a 4 mm diameter punch hole to puncture disks for LMA measurements. We collected one disk per leaf (see Supplementary material for details). The dried disk punches were shipped to Tulane University, New Orleans, LA, USA to dry at 60 ℃ for 48-72 hours before being weighed.

### 3.3 Molecular Work

#### 3.3.1 Tissue preservation

We assessed FEF community composition using an amplicon sequencing approach. Upon completion of the leaf traits measurements, we prepared and preserved samples at the UC Merced Yosemite Field Station. We started by removing the main vein and margins from photosynthetic tissue. The leaf lamina was haphazardly cut with a sterile blade into 2 mm wide strip in parallel to the main vein (Arnold et al., 2003; Higgins et al., 2014; Tellez et al., 2022). Leaf strips were then sterilized with sequential washes in 95% EtOH (10 s), 0.5% sodium hypochlorite (NaOCl) (60 s), and 70% EtOH (60 s) and air dried under sterile conditions. Due to the small size of monkeyflower plants, the maximum amount of leaf lamina was preserved in sterile 15 mL tubes with ∼ 10 mL CTAB solution (1 M Tris–HCl pH 8, 5 M NaCl, 0.5 M EDTA, and 20 g CTAB). The leaf tissue in CTAB solution was used for amplicon sequencing (described in detail below). All leaf tissue handling was performed in a sterile environment with an alcohol burner lamp inside a portable biosafety cabinet. All surfaces were previously sterilized sequentially with 0.5% NaOCl, 95% EtOH, and 70% EtOH. We surface sterilized surfaces and instruments in between sample handling to prevent cross contamination.

#### 3.3.2 Amplicon sequencing

We stored leaf tissue in CTAB solution for 2 months at room temperature before extracting DNA at Tulane University. To prepare for sample DNA extraction procedure, we decontaminated all instruments, materials, and surfaces in biosafety cabinet with 0.5 % NaOCl, 70 % EtOH, and 95% EtOH, and subsequently treated with UV light for 30 minutes. We subsampled 0.2 - 0.3 g of leaf tissue from each sample and placed into a sterile 2 mL tubes containing an assortment of beads: 3.2 mm stainless steel beads (Next Advance, Cat# SSB32), 100 µL stainless steel bead blend, 0.9-2.0mm (NextAdvance, Cat# SSB14B) and 2-3 of the autoclaved 2 mm zirconium oxide beads (Next Advance, Cat# ZRoB20). The 2 mL tubes with beads were previously prepared. We then proceeded to lyophilize samples for 72 hours to fully remove CTAB content from tissue. After, we submerged the sample tubes in liquid nitrogen for 30 s and homogenized samples at 30 Hz for 3 minutes in a TissueLyser LT (QIAGEN, Valencia, CA, USA). We stored samples in 20 ℃ until DNA extraction procedure.

We used a DNA extraction protocol for high-molecular weight DNA extraction adapted from Russo et al., (2022). Briefly, it is a CTAB:chloroform:isoamyl extraction combined with a solid-phase reversible immobilization (SPRI) bead step (Rohland and Reich, 2012; Russo et al., 2022; Liu et al., 2023). Protocol modifications allowed us to optimize extractions for fungal DNA from preserved leaf tissue (see details in Aponte Rolón, 2023). After all genomic DNA was extracted, we quantified the DNA using Quant-iT® dsDNA HS Assay kit with Qubit Flourometer (Thermo Fisher Scientific, Waltham, MA, USA., Cat# Q33120) and followed a two-step amplification approach described by Sarmiento et al. (2017) and U’Ren & Arnold (2017). We used standard primers ITS1F (Gardes and Bruns, 1993) and ITS2 (White, T. J. et al., 1990) modified with the Illumina TruSeq adaptor (see Supplementary Material). Every sample was amplified in three parallel reactions at the annealing temperatures 52 ℃, 54 ℃, and 56 ℃ to amplify a wide range of fungal taxa and reduce amplification bias for short ITS sequences (U’Ren and Arnold, 2017; Lumibao et al., 2018). Each PCR (PCR1) reaction contained 2 µL of sample DNA template. We visualized PCR1 reactions with SYBR™ Safe DNA Gel Stain (Thermo Fisher Scientific, Waltham, MA, USA., Cat# S33102) on 2% agarose gel (Oita et al., 2021). We combined 5 µL of amplicon product from parallel reactions into a single tube per sample and purified using Sera-Mag™ SpeedBead Carboxylate-Modified Magnetic Particles (Hydrophobic) (Thermo Fisher Scientific, Waltham, MA, USA., Cat#09-981-123) prepared as per (2023) and used a ratio of 1.2x:1 with 80% EtOH following manufacturer’s instructions. We used 3 µL of PCR1 product from samples, DNA extraction controls, and PCR1 negative controls for a second PCR (PCR2) with barcoded adapters (IDT, Coralville, Iowa, USA). Each PCR2 reaction (total 30 µL) contained 1X Phusion Flash High Fidelity PCR Master Mix (Thermo Fisher Scientific, Waltham, MA, USA., Cat# F548L), 0.075 µM of barcoded primers (forward and reverse pooled at an initial concentration of 2 µM) and 0.20mg/mL of BSA (Thermo Fisher Scientific, Waltham, MA, USA., Cat# B14) following (2017). Before final pooling for sequencing, we purified and concentrated amplicons using SPRI beads to a total volume of 20 µL. We quantified PCR2 product with Quant-iT™ PicoGreen™ dsDNA Assay Kit (Thermo Fisher Scientific, Waltham, MA, USA., Cat# P7589) with the BioTek Synergy LX plate reader (Agilent, Santa Clara, CA) and combined equimolar amounts of libraries, including DNA extraction controls, PCR1, and PCR2 negative controls into a 10nM library pool. We did not detect any contamination visually or fluorometrically. Libraries were sequenced on the Illumina MiSeq platform with Reagent Kit v3 (2 0D7 300 bp) at Duke Genome Sequencing and Analysis Core Facility (Durham, NC, USA). Throughout all these steps, we used a separate set of sterile pipettes, tips, and equipment to reduce contamination in a designated PCR area to restrict contact with pre-PCR materials (Oita et al., 2021).

#### 3.3.3 Bioinformatic analyses

We assessed the quality of the reads using FastQc v0.12.1 [ v0.12.1; Andrews et al. (2010)] and MultiQC (Ewels et al., 2016) tools. A total of 60,696,808 total ITS1 reads yielded from 343 (including 27 controls) libraries sequenced in two separate sequencing runs. The first sequencing event yielded 32,117,684 and the second 28,579,124 ITS1 reads. We tailored the open-source DADA2 bioinformatic pipeline for our data set (Callahan et al., 2016). Based on our initial quality assessment, both forward and reverse reads were of low quality, with base calls deteriorating after 100 bp. We filtered our reads for ambiguous calls before removing the adapters by using `filterAndTrim` function and argument `maxN = 0` from the `dada2` package [v1.28.0; Callahan et al. (2016)]. We removed forward and reverse primer adapters (and their reverse compliments) and eliminated reads shorter than 20 bp using the `cutadapt` tool (v4.6, Martin, 2011). After removing ambiguous calls and forward and reverse primers, we assessed the quality of the reads again and saw slight improvement. We proceeded to apply stringent filter and truncation parameters to ensure quality of reads when assigning taxonomy. We filtered and truncated reads based on maximum expected errors (maxEE) rather than read length as it provides a reliable quality filtering (Edgar and Flyvbjerg, 2015). For this we set the arguments `trunQ = 2`, `maxEE = c(2,2)` for forward and reverse reads, and minimum read length of 50 bp with `minLen = 50` in the used the `filterAndTrim` function (Callahan et al., 2016). These parameters eliminated 151 samples from our data set, all from our second sequencing event. After this filter, we dereplicated reads with the `derepFastq` function and merged pairs using `mergePairs` functions with an overlap of 20 bp, minimum. We then inferred composition of the samples with `dada` function, which applies the DADA algorithm (Rosen et al., 2012; Callahan et al., 2016). We removed chimeras via the “consensus”method with the `removeBimeraDenovo` function and ultimately we used the `assignTaxonomy` function to assign taxonomy the amplicon sequence variants (ASV) referenced against the UNITE database (Abarenkov et al., 2023). After taxonomy assignment we used the `phyloseq` package (McMurdie and Holmes, 2013) to create a phyloseq object for downstream analyses.

We used the `decontam` package [v1.20.0; Davis et al. (2018)] to statistically determine which ASVs are likely contaminants based on their frequency in our samples and remove them using `prune_taxa` function from the `phyloseq` package [v1.44.0; McMurdie and Holmes (2013)]. After which, we calculated the average read count found in DNA and PCR extraction controls, considered to be laboratory contaminants, and subtracted that from the samples’ read counts. We then used custom scripts to remove any ASV that represented less than 0.1% of the abundance per sample on the assumption that it originates from contamination throughout handling of samples in the DNA and PCR processes. We removed singleton ASVs with the `prune_taxa` function (McMurdie and Holmes, 2013). We identified core taxa members at a 1% detection threshold and 50% prevalence in samples using the `core_members` function from the `microbiome` package [v.1.22.0; Lahti and Shetty (2012–2019)]. All post-quality bioinformatic steps were performed in *R* [v.4.4.1; R Core Team (2024)].

### 3.4 Statistical Analyses

#### 3.4.1 Leaf traits

We checked for normality and homoscedasticity of the leaf traits measured. We used Shapiro-Wilk and Fligner-Killen tests from the `stats` package (R Core Team, 2024) to check for normality and homoscedasticity, respectively. We established that the leaf functional trait data was not normally distributed and not homoscedastic. We then used non-parametric tests, the Wilcoxon Rank Sum test, to compare leaf functional trait means among species and sites to answer the first portion of Q1 and Q2. We used the `compare_means` and `stat_compare_means`functions from the `ggpubr` package (Kassambara, 2023) to perform these tests and properly visualize them. We adjusted *p* values to account for false discovery rates in multiple comparisons by using “BH” method (Benjamini and Hochberg, 1995). We performed Principal Component Analysis (PCA) to understand patterns and relationships among leaf traits of host species. We used the `prcomp` function from the `vegan` package (Oksanen et al., 2022) compute the PCA analysis with variables ACI, LT, LPS, LMA, and LBI, all log-transformed. We used only complete raw leaf functional traits measurements to compute the PCA analysis (*n* = 504), samples with missing values were omitted. All statistical analyses were performed in *R* programming language [v.4.4.1; R Core Team (2024)]. We present the log-transformed leaf functional trait data at the leaf level from a total of 309 plants: ACI (*n* = 851), LT (*n* = 927), LPS (*n* = 875), LMA (*n* = 591), LBI (*n* = 769). The FEF community data is presented from a subset of individuals at the plant/sample level (*n* = 157).

#### 3.4.2 Community Diversity

To account for uneven sampling effort and over-representation of sequences, we normalized libraries by repeated rarefying, following Cameron et al. (2021). We determined a sequence depth of 750 by calculating Good’s coverage and qualitative evaluation of libraries to determine a balanced coverage and breadth of samples (Schloss, 2024). This approach allowed for a proportionate representation of observed sequences from host species and a robust characterization of random variation inherent in rarefaction of small libraries (Cameron et al., 2021; Schloss, 2024). First, we randomly selected 136 samples out of the TRUE viable samples that resulted from the bioinformatic pipeline (refer to the Results section below). The sample pool was reduced to 136 to match the theoretical reduction of one sample per site (*n* = 21). We generated 50 rarefied abundance matrices without replacement by using the `mirl` function from the `mirlyn` package (Cameron et al., 2021). After which we were left with 84 samples. We then calculated α-diversity per sample as Hill orders, the observed richness (*q* = 0), the exponent of Shannon’s entropy (*q* = 1), and the Inverse Simpson’s Diversity (*q* = 2) by applying a modified version of the function `alphadivDF` (Cameron et al., 2021), which wraps the common diversity indices in `vegan` (Oksanen et al., 2022) (see custom script in Supplementary material). Finally, for beta diversity analyses, we performed a Hellinger transformation on the rarefied abundance matrices and calculated a Bray-Curtis dissimilarity matrix for each.

We calculated simple linear regressions and non-parametric tests, the Wilcoxon Rank Sum test, to understand how α-diversity changed in response to elevation (Q1) and (Q2). To facilitate our understanding of the effects of elevation on FEF α-diversity, we categorized elevation as “low” (< 1219 m.a.s.l), “mid” (1220 - 1828 m.a.s.l.) and “high” (> 1829 m.a.s.l.). To statistically compare the FEF community similarities within each host species across sites (Q3) we applied a distance-based Redundancy Analysis (dbRDA) on the Bray-Curtis dissimilarities which effectively describes underlying patterns of compositional differences (Legendre and Anderson, 1999; McArdle and Anderson, 2001; Anderson, 2017). We selected model terms by calculating Spearman’s rho for all leaf traits (see below and Fig. S1) to understand which leaf functional traits were correlated with FEF richness, diversity and community composition. We informed our selection of leaf traits with results from the PCA (see below and Fig. 1), and selected those with the lowest correlation coefficient per host species: logLBI. Our initial dbRDA model consisted of terms logLBI, sampling date, site, and elevation (m). The leaf functional trait data, as well as elevation, was not randomized or subsampled to match rarefied dataset, the same values applied to all 50 rarefied matrices. For our initial model, we determined the variance inflation factor (VIF) of each term with function `vif.cca` and eliminated redundant ones: site and sampling date. We performed manual model selection by evaluating the marginal significance of constraining variables after 999 permutations with function `anova.cca` and argument `by` set to “terms” to assess significance for each term (Legendre et al., 2011; Legendre and Legendre, 2012; Oksanen et al., 2022). We corroborated homogeneous dispersion of variances with a permutational analysis of multivariate dispersion (PERMDISP) using the `betadisper` with parameter `type = “median”`, and `permutest` functions from `vegan`, the latter with 999 permutations (Oksanen et al., 2022). We used a post-hoc Tukey’s test to compare the differences in the dispersion of the FEF communities among sites and elevation zones. We used the `simper` function from `vegan` to discriminate which species contribute the most to compositional differences between groups (Oksanen et al., 2022).

We used generalized linear mixed models (GLMM) to determine the effect of host species, elevation, and leaf functional traits on the mean Bray-Curtis dissimilarity (β-diversity) of FEF communities (Q3). Leaf functional traits with missing values were imputed with the median value for the trait. We used the `lme` function from the `nlme` package [v3.1-165; Pinheiro et al. (2024); Pinheiro and Bates (2000)] to fit the GLMM models. We established leaf functional traits, elevation and host species as fixed terms and site as a random effect. We modeled variance structure for site with the `varIdent` argument. We manually compared and selected models based on Akaike Information Criterion (AIC) with a penalty of 2 degrees of freedom (ΔAIC) with the `model.sel` function from the `MuMIn` package (Bartoń, 2023). We selected the best-fit model based on the lowest value obtained. All models are based on restricted maximum likelihood estimation (REML).

To assess correlations between pairwise FEF community dissimilarity and the geographical distance matrix per host species (Q4), we computed Mantel tests with Spearman’s rho and 999 permutations using the `mantel` function (Oksanen et al., 2022). For this test, we opted for a less taxing computational approach and rarefied sequences with the same parameters as before and calculated Hellinger transformations with the `avgdist` function (Oksanen et al., 2022). We then calculated the Bray-Curtis dissimilarity with `vegdist` (Oksanen et al., 2022). For the geographical distances between sites, we used `distm` function with the Vicenty (ellipsoid) method from the `geosphere` package (Hijmans, 2022).

## 4. Results

### 4.1 Leaf traits differ among host species and separate *M. laciniatus* via lobing

We found evidence of both interspecific (Q1) and intraspecific elevational shifts (Q2) in leaf functional traits among sympatric *Mimulus* species (Fig. 2 and Figs. S5–S9). Our PCA analysis showed how leaf functional traits varied among sympatric *Mimulus* species populations at similar elevations (Q1). We plotted leaf functional traits according to species groups on the PCA axes to show how the variance in the complete data set is explained by PC1 (42.57%) and PC2 (23.27%) (Fig. 2). The PCA showed correlations between ACI, LT, LPS, and LMA as loadings that tracked along PC1 towards positive values (Fig. 2). We observed that LBI loading was orthogonal along PC2 to the other traits, indicative of low correlation. We noted distinct groupings by species along PC2 such that *M. laciniatus*, the most lobed species, was distinct from *M.guttatus* and *M. nasutus*. The latter two overlapped along PC1 and PC2 (Fig. 2).

**Figure 2:**
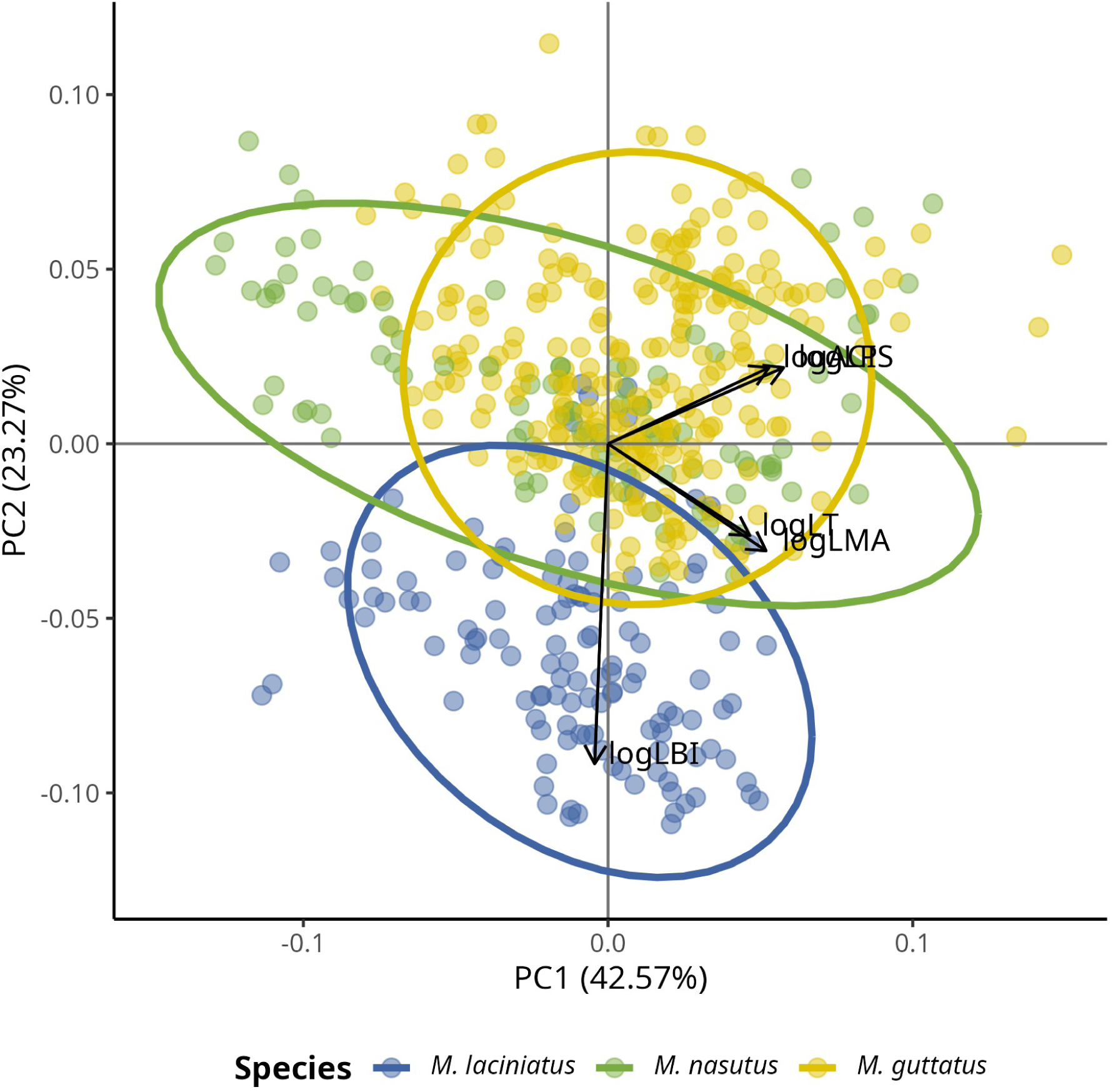
The Principal Component Analysis (PCA) shows how leaf functional traits are correlated. The PCA analyses shows correlations between ACI, LT, LPS, and LMA as loadings track along PC1 towards positive values, while leaf lobing correlates positively with M. laciniatus. We observed distinct groupings by species along PC2 such that *M. laciniatus*, the most lobed species, is distinct from *M. guttatus* and *M. nasutus*. The latter two have greater overlap along PC1 and PC2.

#### 4.1.1 Host species differ in leaf traits at shared elevations

At shared elevations, leaf functional traits frequently showed interspecific differences, though the magnitude and direction of differences varied across elevation zones (Figs. S5B–S9B). At mid-elevations, where all three species co-occurred, *M. laciniatus* exhibited distinct trait values compared to *M. guttatus* and *M. nasutus*, including higher leaf mass per area (LMA, *p* < .001; Fig. S5B), lower anthocyanin content (ACI, *p* < .0001; Fig. S6B), and greater lobing (LBI, *p* < .0001; Fig. S7B). At high elevations, *M. nasutus* diverged from both congeners in LMA (*p* < .0001; Fig. S5B) and ACI (*p* < .0001; Fig. S6B), while *M. guttatus* and *M. nasutus* differed in leaf thickness (LT, *p* < .0001; Fig. S9B). Notably, LBI showed minimal overlap among species across all elevations (Fig. S7B), reflecting strong species-specific morphological constraints.

#### 4.1.2 Trait–elevation relationships are species-specific

Trait-elevation relationships varied by species and trait type (Q2). Across the entire community, LMA, LPS, and LT increased weakly but significantly with elevation (LMA: *R²_adj_* = .041, *p* < .001; LPS: *R²_adj_* = .004, *p* = .032; LT:*R²_adj_* = .013, *p* < .0001; Figs. S5A, S8A, S9A). However, categorical analyses revealed species-specific patterns: *M. laciniatus* maintained consistently high LBI across elevations (Fig. S7B), while *M. guttatus* showed increased LPS at mid-elevations (*p* < .0001; Fig. S8B) and greater LT differentiation at higher elevations (*p* < .0001; Fig. S9B). In contrast, ACI showed no community-wide elevational trend (*R²_adj_* < −.000, *p* < .0001; Fig. S6A), but *M. laciniatus* exhibited lower ACI at higher elevations compared to congeners (*p* < .01; Fig. S6B).

#### 4.1.3 FEF communities shift with elevation and geography

##### 4.1.3.1 FEF communities are Basidiomycete-dominated across hosts and elevations

We obtained 5,082,229 reads representing 726 ASVs from 174 samples after processing samples through the DADA2 pipeline. The raw reads obtained were composed of 26.81% Ascomycota, 71.53% Basidiomycota, <0.05% Chytridiomycota, <0.5% Mortierellomycota, <0.03% Olpidiomycota, 0.01% Rozellomycota, <0.001% Aphelidiomycota, and 1.19% unidentified. After decontamination, and removal of singletons we retained 4,856,220 reads representing 231 ASVs from 157 samples composed of 26% Ascomycota, 73% Basidiomycota, 0.01% Chytridiomycota, <0.1% Mortierellomycota, 0.03% Olpidiomycota, <0.002% Rozellomycota and <1.0% unknown reads (Fig. 3 and Fig. S2). After rarefaction of sequences, we were left with 84 samples in which we found the most prevalent taxa: *Vishniacozyma victoriae* (Basidiomycota, ASV_1), *Cladosporium herbanum* (Ascomycota, ASV_2) and *Cladosporium* spp. (Ascomycota, ASV_7), *Dyszogia patagonica* (ASV_3), *Filobasidium chernovii* (Basidiomycota, ASV_5), and *Alternaria tenuissima* (Ascomycota, ASV_8) (Fig. S2).

**Figure 3:**
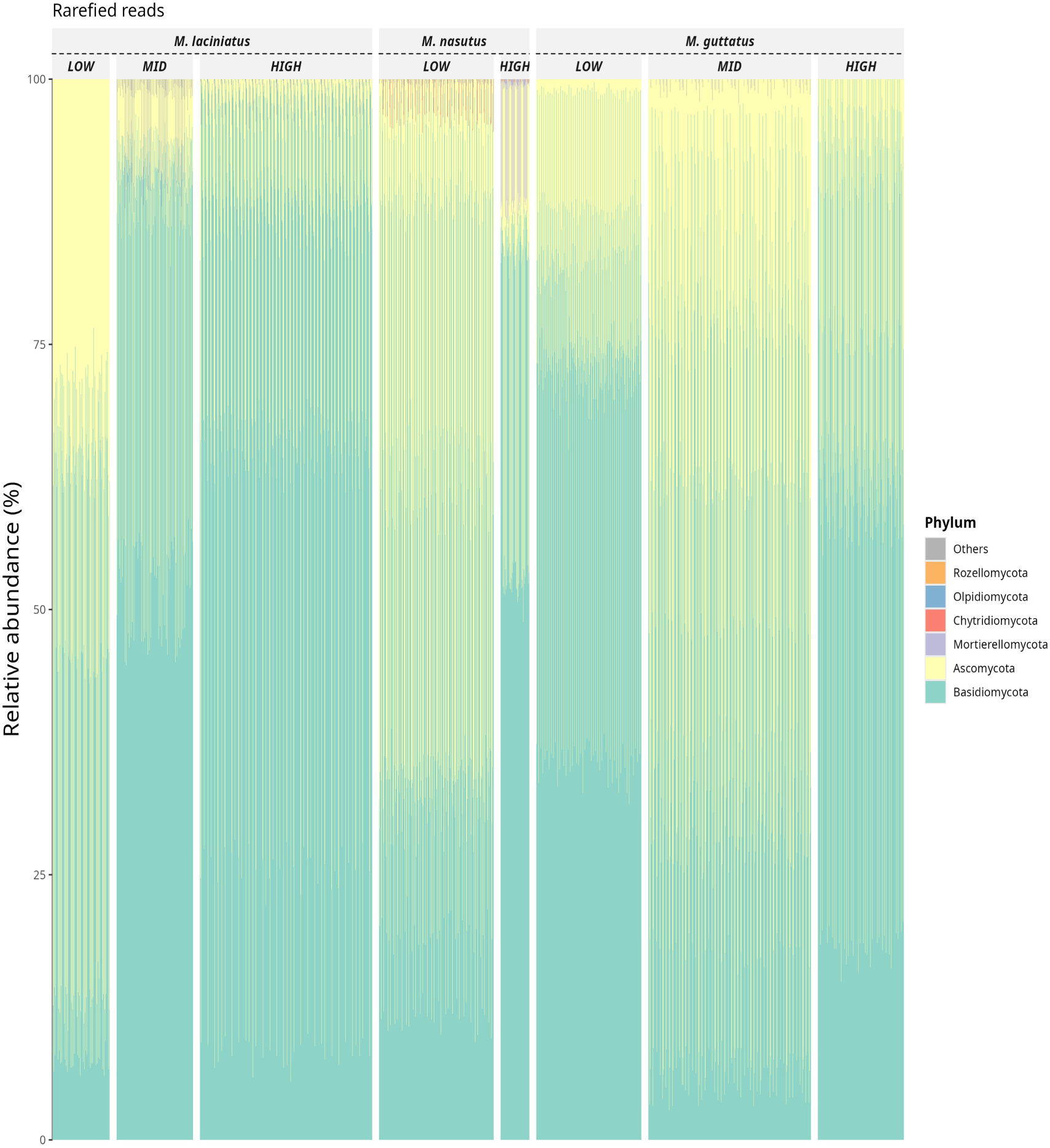
Relative abundance of fungal phyla in rarefied sequence data by species and elevation zone.

#### 4.1.4 Elevation and host species shape FEF diversity and composition: leaf traits predict β-diversity

##### 4.1.4.0.1 FEF α-diversity declines with elevation, but responses are species specific

We observed significant interspecific differences in α-diversity across all Hill numbers (*q* = 0, 1, 2), with the sole exception of the *M. laciniatus* – *M. nasutus* contrast at *q* = 0 (Fig. S4A–S4C) (Q1 and Q3). FEF α-diversity decreased significantly with elevation across all Hill numbers: (*q* = 0) (*R^2^* < .01, *F* = 16.69, *p* < .001), the exponent of Shannon’s entropy (*q* = 1) (*R^2^* < .01, *F* = 37.91, *p* < .001), and the Inverse Simpson’s Diversity (*q* = 2) (*R^2^* < .01, *F* = 23.73, *p* < .001). Species-specific responses differed: *M. nasutus* showed declining diversity with elevation (Fig. 4), while *M. laciniatus* exhibited increased diversity. *M. guttatus* displayed mixed trends, with elevated Hill *q* = 0 diversity but decreased diversity at higher Hill orders. We observed a similar pattern in beta diversity among elevation sites (Fig. S4D -S4F). We saw no differences in beta diversity between low and mid elevation sites for Hill order 2 (*q* = 2, *p* > .05, Fig. S4F).

**Figure 4:**
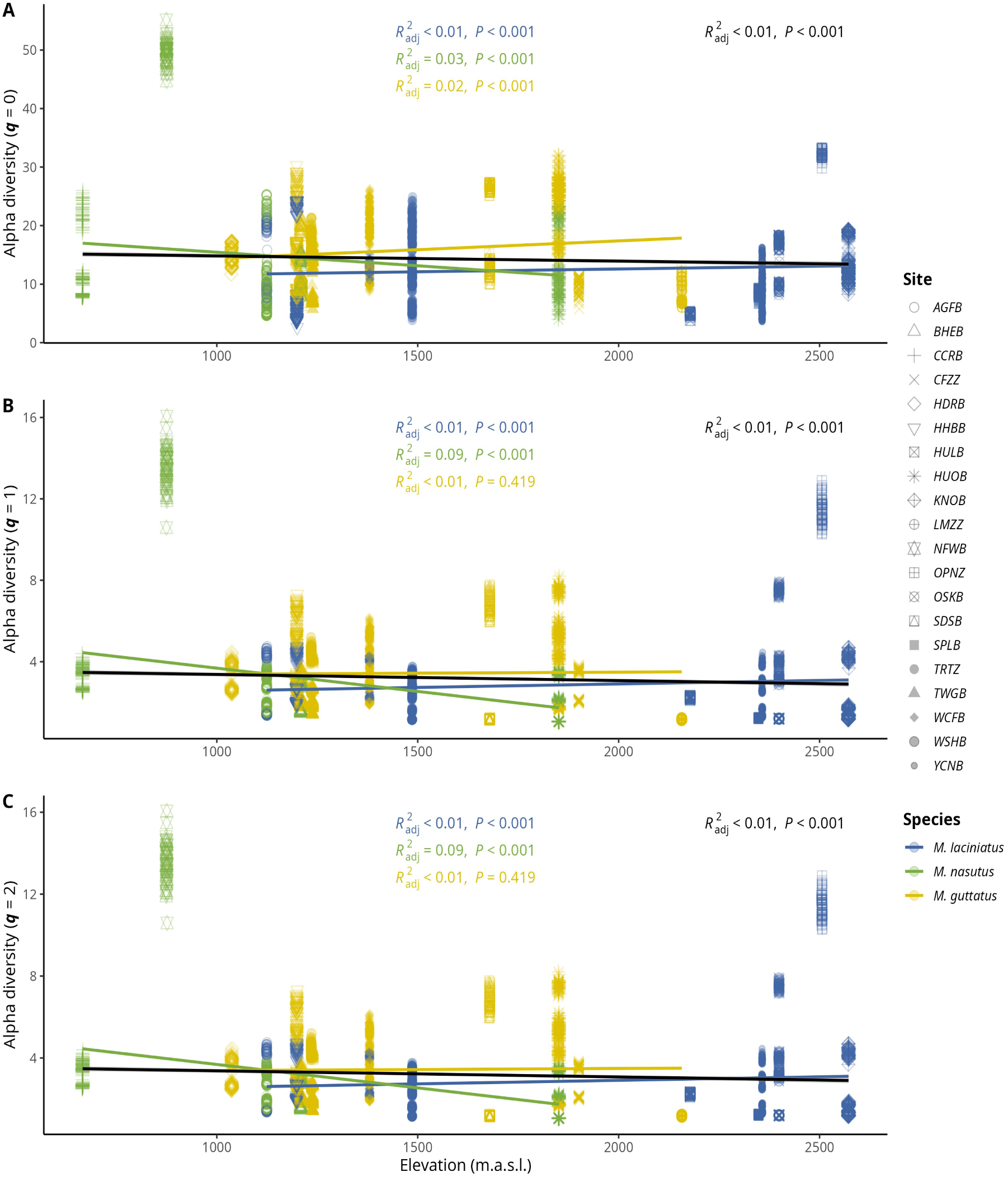
Relationship between Hill orders by host species and elevation. A) Observed ASV richness (*q* = 0); B) Shannon’s entropy (*q* = 1); and C) Inverse Simpson’s index (*q* = 2) per host species as elevation increases. Blue filled points correspond to *M. laciniatus*, green filled to *M. nasutus*, and yellow to *M. guttatus*. The solid line represents the linear model fit and the shaded area represents the 95% confidence interval.

##### 4.1.4.0.2 Host species and elevation jointly structure FEF community composition

FEF communities differed significantly among host species and elevation zones. The best-fit model revealed that 22% of the overall variance in FEF communities was accounted for by log-transformed LBI, and the interaction of host species and elevation, all of which were significantly correlated (*p* < .001) with FEF communities in their host species (Fig. 5 and Table S3). We observed that the first axis (dbRDA1) explained 51.7% and the second axis (dbRDA2) explained 15.9% of the constrained variance (Fig. 5) which support that host species and elevation interactions structure FEF communities. Communities showed greater overlap at low elevations but became distinct at high elevations (Fig. 5).

**Figure 5:**
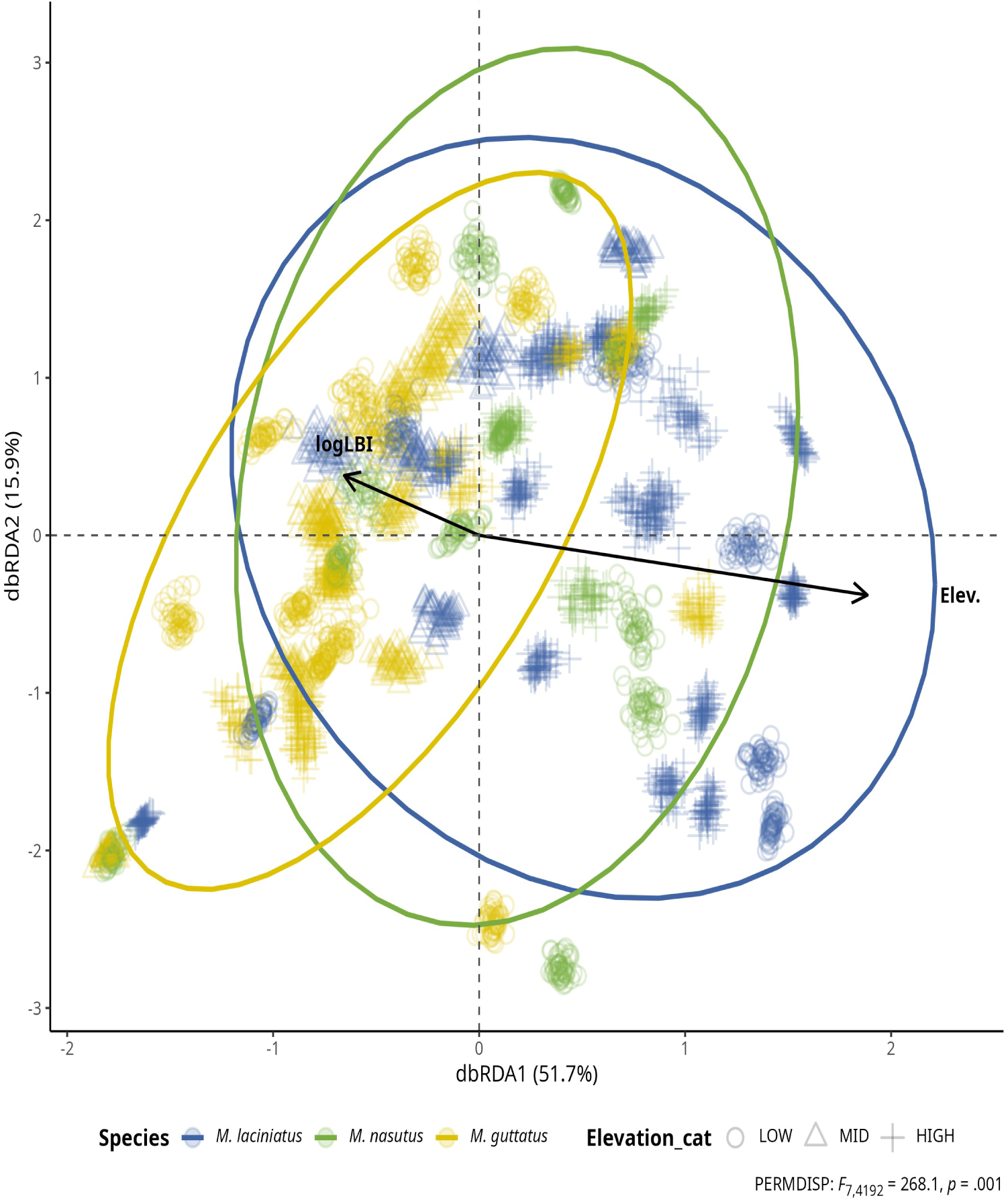
FEF community composition association with leaf functional traits and elevation by host species. Distance-based Redundancy Analysis (dbRDA) plot of rarefied FEF community and leaf functional traits by species (colors) and by elevation zone (shapes). Each cluster of points represents a rarefied FEF community sample from one host species. Solid arrow lines represent significant associations (*p* < .01) with non-significant leaf trait associations left out for clarity. The length and direction of the arrows indicate the strength and direction of the association between the traits and the FEF community composition. Ellipses represent 95% confidence intervals. The plot is based on the Bray-Curtis dissimilarity matrix.

We found that FEF communities were not homogeneous across all species (*F*_2, 4197_= 320.3, *p* < .001). The post-hoc Tukey’s test revealed that all species comparisons were significantly different at ⍺ = 0.05. We detected significant differences in the dispersion of FEF communities by elevation zone (*F*_2, 4197_= 228.1, *p* < .001) (Tukey’s test ⍺ = 0.05). The interaction between log LBI and elevation zone also showed differences in group dispersion (*F*_2, 4192_= 268.1, *p* < .001). Nevertheless, PERMDISP tests indicated significant dispersion differences for most comparisons across all species (*F*_2, 4197_= 320.3, *p* < .001), elevation zones (*F*_2, 4197_= 228.1, *p* < .001) and the interaction between log LBI and elevation zone (*F*_2, 4192_= 268.1, *p* < .001) (Tukey’s test ⍺ = 0.05). Notably, dispersion patterns were homogeneous for: *M. guttatus* across all elevations, *M. laciniatus* (LOW/MID) vs. *M. guttatus* (HIGH), and *M. nasutus* (MID/HIGH) vs. *M. laciniatus* (MID) (Tukey’s test, ⍺ = 0.05). While we cannot rule out that the observed differences FEF communities are due to dispersion, they are unlikely to fully explain the observed community structure.

##### 4.1.4.0.3 Leaf functional traits, not elevation, predict β-diversity

GLMMs identified leaf functional traits, not elevation, as primary predictors of mean β-diversity (Model 4, Table 1). Anthocyanin content (ACI) showed the strongest negative association (estimate = −0.078, *t*(4200) = −32.48, *p* < .001), followed by leaf thickness (LT; estimate = −0.051, *t*(4200) = −9.51, *p* < .001) and leaf mass per area (LMA; estimate = 0.013, *t*(4200) = 3.74, *p* < .001). Simple linear regressions showed that *M. nasutus* (*R^2^* = .46, *F* = 642.3, *p* < .001) and *M. guttatus* (*R^2^* < .01, *F* = 38.3, *p* < .001) exhibited elevational declines in β-diversity, while *M. laciniatus* showed no elevational trend (*R^2^* < .01, *F* = 1.84, *p* = .176, Fig. 6).

**Figure 6:**
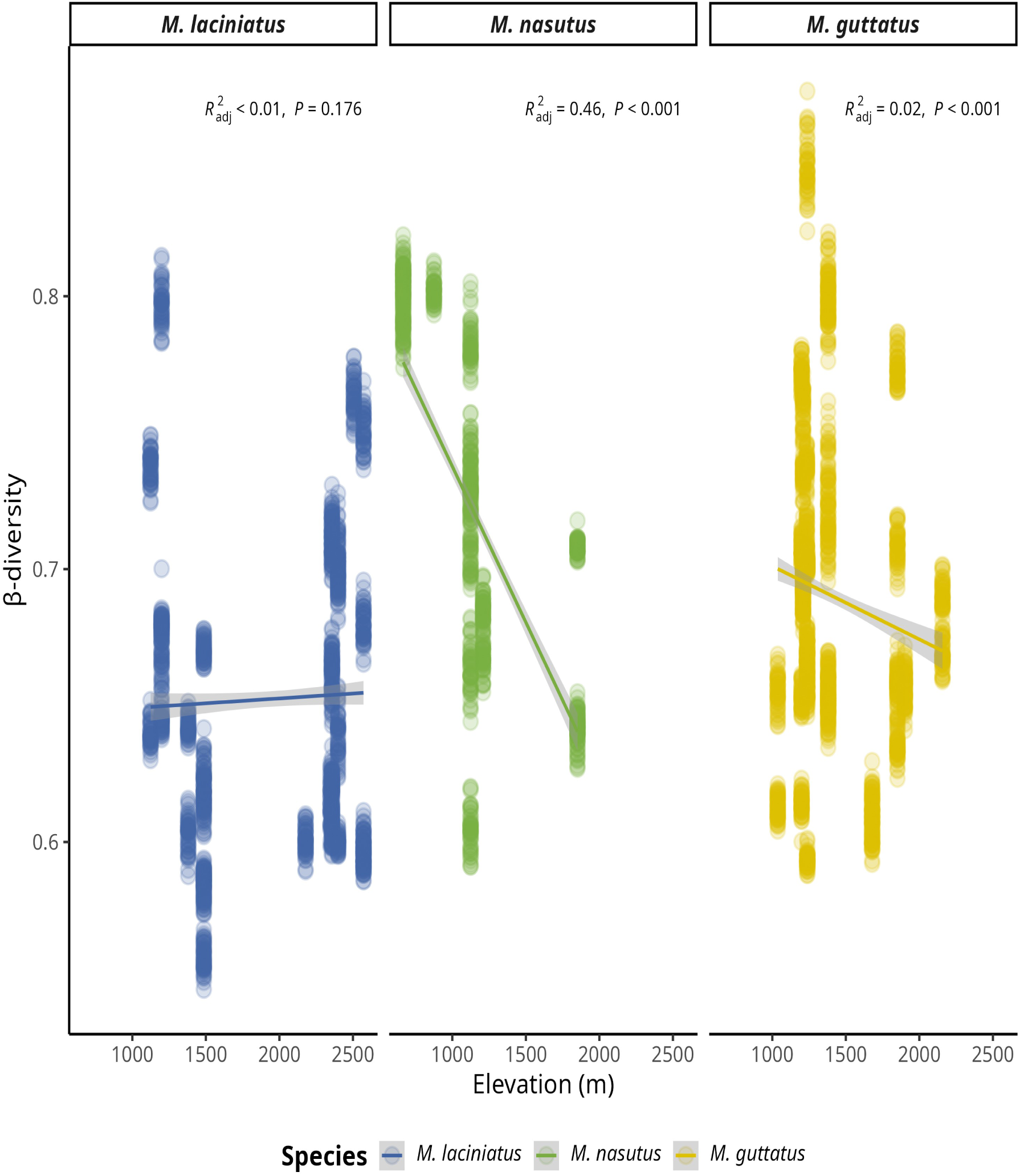
Relationship between β-diversity and elevation by host species. The solid line represents simple linear regression and the shaded area represents the 95% confidence interval.

#### 4.1.5 Geographic distance predicts FEF turnover in M. laciniatus and M. nasutus but not M. guttatus

Finally (Q4), we detected clear distance–decay in FEF mean β-diversity turnover in *M. laciniatus* (*r* = 0.27, *p* = .01) and *M. nasutus* (*r* = 0.29, *p* < .01, Fig. 7), consistent with isolation by distance and/or spatially structured environmental filters acting on foliar fungi. In contrast, *M. guttatus* (*r* = −0.004, *p* = .48) showed no evidence of distance-decay (Fig. 7), consistent with broader dispersal of its dominant endophytes, greater host gene flow that weakens population structure and geographic filtering, or stronger local environmental/host-trait filtering that overwhelms spatial effects. Distance–decay in two selfing species but not in *M. guttatus* suggests that dispersal limitation and host spatial structure shape FEF communities, with species-specific balances between geographic and environmental filtering.

**Figure 7:**
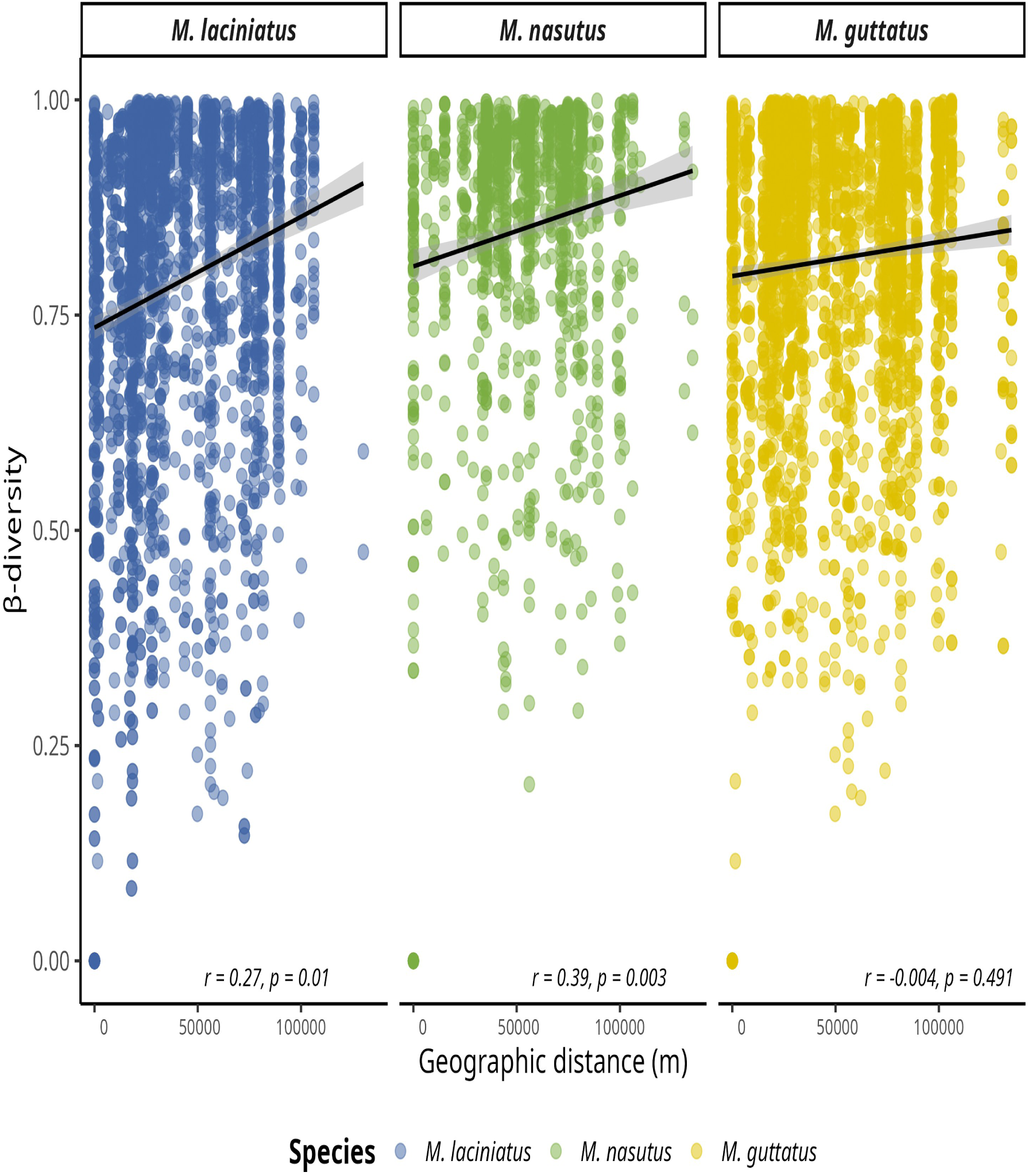
Dissimilarity of FEF communities correlates with *Mimulus* as a function of geographic distance per site. Data represent pairwise Bray-Curtis dissimilarity (β-diversity) among sites. A) FEF community dissimilarity in considering all host species. B) FEF community dissimilarity by host species. Blue filled points correspond to *M. laciniatus*, green filled to *M. nasutus*, and yellow to *M. guttatus*.

## 5. Discussion

In this study, we examined how leaf functional traits and foliar endophytic fungal (FEF) communities vary among three sympatric *Mimulus* species across elevation gradients, and whether elevation, host identity, and leaf traits jointly structure FEF richness, diversity, composition, and spatial turnover. Below, we first discuss inter- and intraspecific patterns in leaf functional traits along elevation, then interpret elevational trends in FEF alpha and beta diversity, followed by trait/elevation effects on community composition and dispersion, geographic distance–decay patterns within species, and finally the consistent dominant taxa observed across sites, along with methodological considerations and future directions.

### 5.1 Closely related sympatric Mimulus differ in leaf functional traits

We found clear interspecific differentiation in leaf functional traits among sympatric *Mimulus* species, with leaf lobing (LBI) providing a particularly strong axis of separation among hosts. Specifically, LMA, LBI, LPS and LT, were positively correlated with elevation at the genus level, reflecting a conserved response to environmental changes along elevation. The Principal Component Analysis (PCA) (Fig. 2) illustrated correlations between leaf functional traits while demonstrating the distinct groupings of *M. laciniatus*, *M. guttatus*, and *M. nasutus* based on LBI differences compared to the rest of the traits: ACI, LMA, LPS and LT. *Mimulus laciniatus* is the most lobed species (Fig. 2 and Fig. S7) (Sexton et al., 2013; Ferris et al., 2015; Ferris, 2016; Tataru et al., 2023), and thrives at higher elevation. For serrated/lobed leaves, it has been observed that physiological adaptations in the leaf margin are a response to the average air temperature, but this is not a simple relationship (Nicotra et al., 2011; Tsukaya, 2018).

The elevational increase in leaf lobing and other structural traits suggests that *Mimulus* leaf morphology may track abiotic constraints such as temperature and water availability along the gradient. The observed increase in LBI with elevation, which has been previously documented in *M. laciniatus* by Love and Ferris (2024), could be a response to the need to increase hydraulic efficiency and reduce heat stress in *M. laciniatus* and *M. nasutus*’ drier habitats (Wu et al., 2008; Ferris et al., 2014). Lobed leaves are expected to have increased hydraulic efficiency and have an advantage in arid environments by influencing the leaf boundary layer, an area of still air adjacent the leaf’s abaxial surface (Nobel, 2009; Oguchi et al., 2018). The size of the boundary layer impacts a leaf’s gas and heat exchange (Schuepp, 1993; Nobel, 2009) affecting its response to changes in mean daily temperature and water availability. The increase in LMA, LT and LPS with elevation could demonstrate the need to increase leaf structural strength and reduce water loss at higher elevations in all three species.

Disentangling environmental and genetic drivers of leaf trait variation will require various experimental approaches, particularly common gardens (as in Ferris and Willis (2018)) to fully answer questions about the functional significance of leaf traits for local adaptation to different elevations. Experimental manipulation with *Mimulus* is relatively easily due to its short generation time, high fecundity, self-compatibility, and ease of greenhouse propagation (Wu et al., 2008; Society, 2019). Because these traits can act as host filters on microbial colonization, we next consider whether FEF diversity shows parallel elevational patterns within species.

### 5.2 Within-species symbiont α-diversity declines with elevation

Foliar endophytic fungi α-diversity declined with elevation (Fig. 4), consistent with broader macroecological patterns of biodiversity (Kraft et al., 2011; Sabatini et al., 2018; Villacampa et al., 2019; Jiménez-Hernández et al., 2020), but the strength and direction of this pattern differed among host species. Alpha diversity declined with elevation for *M. nasutus*, *M. laciniatus* exhibited the opposite, while *M. guttatus* displayed mixed trends (Fig. 4). The observed decline in α-diversity appears to be driven by *M. nasutus’* decrease and *M. guttatus*’ unchanged diversity levels with increased elevation. This species specificity motivates examining not only diversity patterns but also how elevation, host identity, and leaf traits structure the composition of FEF communities.

### 5.3 Host species and elevation jointly structure FEF community composition; leaf traits predict β-diversity

Consistent with the idea that both abiotic gradients and host identity filter symbiont communities, we found that host species, elevation, and LBI were associated with differences in FEF community composition. Our dbRDA results (Fig. 4) indicate that host species, elevation, and LBI account for 17% of the variance in FEF community composition. Subsequent PERMDISP analyses also support significant differences in FEF community composition between host species at low, mid, and high elevations. However, when we modeled mean β-diversity directly, leaf functional traits rather than elevation emerged as the strongest predictors, prompting a closer look at how analytical approaches shape inference.

Although ordination-based results highlighted elevation and host identity, GLMM results showed that ACI, LT, and LMA are more direct predictors of mean β-diversity than elevation *per se* (Fig. 6). Apparent discrepancies likely arise from model specification: the GLMM explicitly modeled site-level variance (Table (**tab-table1?**)), whereas the dbRDA did not partition site from elevation, potentially introducing unaccounted site effects.

### 5.4 Geographic distance predicts FEF turnover in M. laciniatus and M. nasutus

Within species, spatial structure also contributed to FEF turnover: community dissimilarity increased with geographic distance in *M. laciniatus* and *M. nasutus* according to our Mantel tests, providing evidence in support of spatially driven differences in FEF community composition. It is interesting to note that the most dissimilar FEF communities due to geographical distance are in the two species that have self-fertilizing mating systems, *M. laciniatus* (Ferris et al., 2014) and *M. nasutus* (Brandvain et al., 2014). This could indicate that mating systems and host species’ genetic structure could play a role in structuring FEF communities. Self-fertilizing species tend to have reduced within population genetic diversity and increased between population divergence due to high levels of inbreeding and reduced gene flow (Sweigart and Willis, 2003; Sexton et al., 2016; Hartfield et al., 2017). Our findings overall support the idea that distinct FEF communities are structured by the interplay of host specific leaf functional traits and geographic distance. To place these patterns in a broader microbial-ecology context, we next compare our results with elevation-gradient studies in other plant systems and tissue types.

Comparisons with other elevation-gradient studies suggest that the relative importance of elevation, host identity, and tissue type can vary substantially across systems. Patterns of diversity and community composition in microbial ecology are often constrained by both biotic and abiotic factors. For example, Bowman and Arnold (2021) showed distance decay in ectomychorrizal fungi (EMF) and foliar endophytes, but drivers differed for each: EMF were constrained by dispersal limitation, whereas foliar endophytes were constrained environmental conditions. Meanwhile, in an experimental setup, Kivlin et al., (2022) reported that host species (alpine grasses) was a stronger predictor than elevation of α-diversity and community composition of leaf endophytes. In contrast, root endophyte communities responded to both host species and elevation (Kivlin et al., 2022). It is possible that our results differ due to the different phenologies and tissue types of herbaceous and gramineous plants. Similarly, Kezenel et al. (2019) found greater change in leaf endophytes due to altitude and warming when compared to root colonizing fungi, but the direction and magnitude of responses varied among host species and fungal functional groups. A major difference in this study is the low ASV count and the use of multiple rarefied data sets compared to Kazenel et al. (2019). A study by Cordier et al. (2012) focused on the fungal phyllosphere in European beech along an elevation gradient, and found that climatic variables, especially temperature, showed the strongest correlations with fungal community dissimilarities. While the phyllosphere of beech varies widely, Cordier et al. (2012) also found a strong affinity of fungal taxa to elevation and site, supporting regional spatial structure. An important distinction is that they focused on the outer and inner phyllosphere, hence observing patterns that reflected the outer leaf dynamics. These may have been more susceptible to climatic factors, as opposed to inner leaf dynamics that we explored in our study. Other key differences are the host species phenology and functional leaf traits of European beech compared to herbaceous *Mimulus* spp.

### 5.5 Consistent taxa underpin FEF communities across elevations

Despite turnover with elevation and distance, several taxa were consistently prevalent across sites and hosts, suggesting a core set of widespread foliar symbionts associated with in Sierran *Mimulus*. The widespread detection of *V. victoriae* across the elevation gradient is particularly notable given its documented associations with cold environments and its proposed functional roles. Additionally, we detected the consistent prevalence of *Cladosporium herbanum*, and *Cladosporium* in across all sites and samples. The basidiomycetous yeast *V. victoriae* (formerly *Cryptococcus victoriae*) is also a well-known environmentally abundant fungus capable of causing respiratory issues in humans (Rush et al., 2023). It was first isolated in the Antarctic (Montes et al., 1999) and has since been detected worldwide (De Menezes et al., 2019). It is proposed that through the production of various bio-active compounds, *V.victoriae* can contribute to plant growth and ecological adaptation to cold environments (Buzzini et al., 2018; Vujanovic, 2021). Despite potential respiratory health detriments, *V. victoriae* has been utilized in agricultural settings for the post-harvest control of fruit diseases (Lutz et al., 2020). It thrives at low temperatures (15 °C), but it is known to tolerate a variety of environmental conditions, and lacks a polysaccharide capsule, which is thought to contribute to its lack of pathogenicity (Rush et al., 2023). Its applications in wheat agriculture suggest that kernel weight is influenced by *V. victoriae*’s coexistence with other plant acquired endophytic fungi (Vujanovic, 2021). Its presence might serve as an indicator of wheat’s kernel resistance to pathobiota (Lutz et al., 2020). We need further quantitative studies to confirm the existence of cold-adapted microbial taxa and their associated hosts (Marian et al., 2022). The presence of *V. victoriae* in our samples might be indicative of its potential role in the local adaptation of *Mimulus* to cold and high elevation environments.

### 5.6 Methodological considerations and future directions

To optimize microbial studies in *Mimulus*, it is crucial to explore alternative sampling methods such as using fresh tissue or rapidly preserved tissue in liquid nitrogen to improve DNA extraction yields. Given the joint signatures of host traits and geography, experiments manipulating host genotype, functional traits, and microbial communities will be essential to test causality. Future studies should consider the role of plant genotype and genetic loci in shaping FEF communities and how these communities might contribute to the host’s adaptation to cold environments.

## 6. Conclusions

Overall, our results show variation in leaf functional traits and FEF diversity, composition, and spatial turnover across elevation in three sympatric Mimulus species. From our results, PCA and trait–elevation trends highlight host filtering; dbRDA shos host–elevation interactions; GLMMs point to leaf functional traits as strong predirctor over elevation for mean β-diversity; Mantel tests reveal distance–decay where dispersal and host structure limit mixing.

Across mountain landscapes, foliar symbiont communities emerge from the joint action of three broad ecological filters: host functional traits, abiotic gradients, and spatial processes. In our system, trait differences among closely related hosts (Q1–Q2), a community-wide decline in α-diversity with elevation (Q3), trait-linked shifts in β-diversity (Q3), and distance–decay in selfing species (Q4) collectively point to an overarching assembly framework in which: (1) Host trait filtering (e.g., leaf thickness, toughness, lobing) sets the local rules of colonization and persistence within leaves; (2)Environmental gradients (e.g., temperature, water availability indexed by elevation) modulate trait distributions and community overlap across sites; (3) Spatial processes (e.g., dispersal limitation, isolation by distance) and host population structure determine the scale and strength of symbiont turnover.

These findings motivate several general ecological questions that extend beyond the focal hosts: (a) What is the relative importance of host trait filtering versus abiotic filtering and how does it shift across spatial scales, plant tissues, and seasons?; (b) How and when does host genetic structure amplify geographic filtering of symbionts? (c) What functional roles do cosmopolitan, cold-associated taxa play in plant performance across climate gradients, and how stable are these roles over time?; (d) How can trait-based and spatially explicit models be integrated to forecast microbiome assembly and function under climate change? (e) To what extent are these assembly rules conserved across plant lineages and fungal guilds, and where do they break down?

Addressing these questions will require synthesis across comparative field surveys, manipulative experiments, and trait-informed community models. In particular, common-garden and reciprocal-transplant designs that jointly manipulate host genotype, leaf traits, and environment can test causal links among filters, while improved tissue preservation and sequencing workflows will sharpen inference on core versus transient taxa. By situating foliar symbionts within a trait–environment–space framework, future work can move from pattern to mechanistic models taht quantify how plant traits and geography govern symbiont assembly, and how those communities feed back to influence plant resilience as climates change and distributions shift.

## 7. Author Contributions

B.A.R, K.G.F. and S.A.V. designed research study. B.A.R. conducted field surveys, laboratory work, and collected data. B.A.R. processed the data and performed statistical analyses. B.A.R. wrote the manuscript with input from all authors. All authors gave final approval for publication.

## 8. Acknowledgements

This work was supported by the National Institute of General Medicinal Sciences of the National Institute of Health (NIH) under award number R35GM138224 to KGF. We thank the Yosemite National Park Service for permit support (YOSE-2021-SCI-0033, YOSE-2022-SCI-0051), and Breeanne Jackson at Yosemite Field Station (doi:10.21973/n3v36c) for logistical support and providing accommodation. We thank the USDA Forest Service for providing access to the Sierra National Forest and Stanislaus National Forest (Botanical Collection Permit: 031528). We thank Caroline, M. Dong, Fidel J. Machado Perez, Elizabeth MacDougal, Rachael Dennis, and Lissette Montano for field assistance. This research was supported in part using high performance computing (HPC) resources and services provided by Information Technology at Tulane University, New Orleans, LA. Lastly, we would like to acknowledge the original inhabitants of the unceded land on which our research was conducted, the Southern Sierra Miwuk Nation, Bishop Paiute Tribe, Bridgeport Indian Colony, Mono Lake Kutzadikaa, North Fork Rancheria of Mono Indians of California, Picayune Rancheria of the Chukchansi Indians and the Tuolumne Band of Me-Wuk Indians. Their culture and stewardship remain an integral part of the land. As scientists we strive to take responsibility for the impacts of colonialism in our field and move forward with respect and support of indigenous movements and knowledge.

## 10. Data Availability Statement

The genomic data that support the findings of this study will be submitted to NCBI-GenBank upon acceptance of this manuscript. The manuscript and code for reproducibility is available from corresponding author’s GitHub:https://github.com/jibarozzo/endophyte_mimulus.git

