## Supplemental_Materials for "Foliar fungal symbionts in sympatric yellow monkeyflowers along elevation gradients in the Sierra Nevada"

January 18, 2026

[1] Department of Agricultural and Biosystems Engineering, Iowa State University, Ames, IA 50011, USA

[2] Department of Ecology and Evolutionary Biology, Tulane University, 6823 St. Charles Avenue, New Orleans, LA 70118

### 13. Supplementary Material

#### 13.1 Figure S1

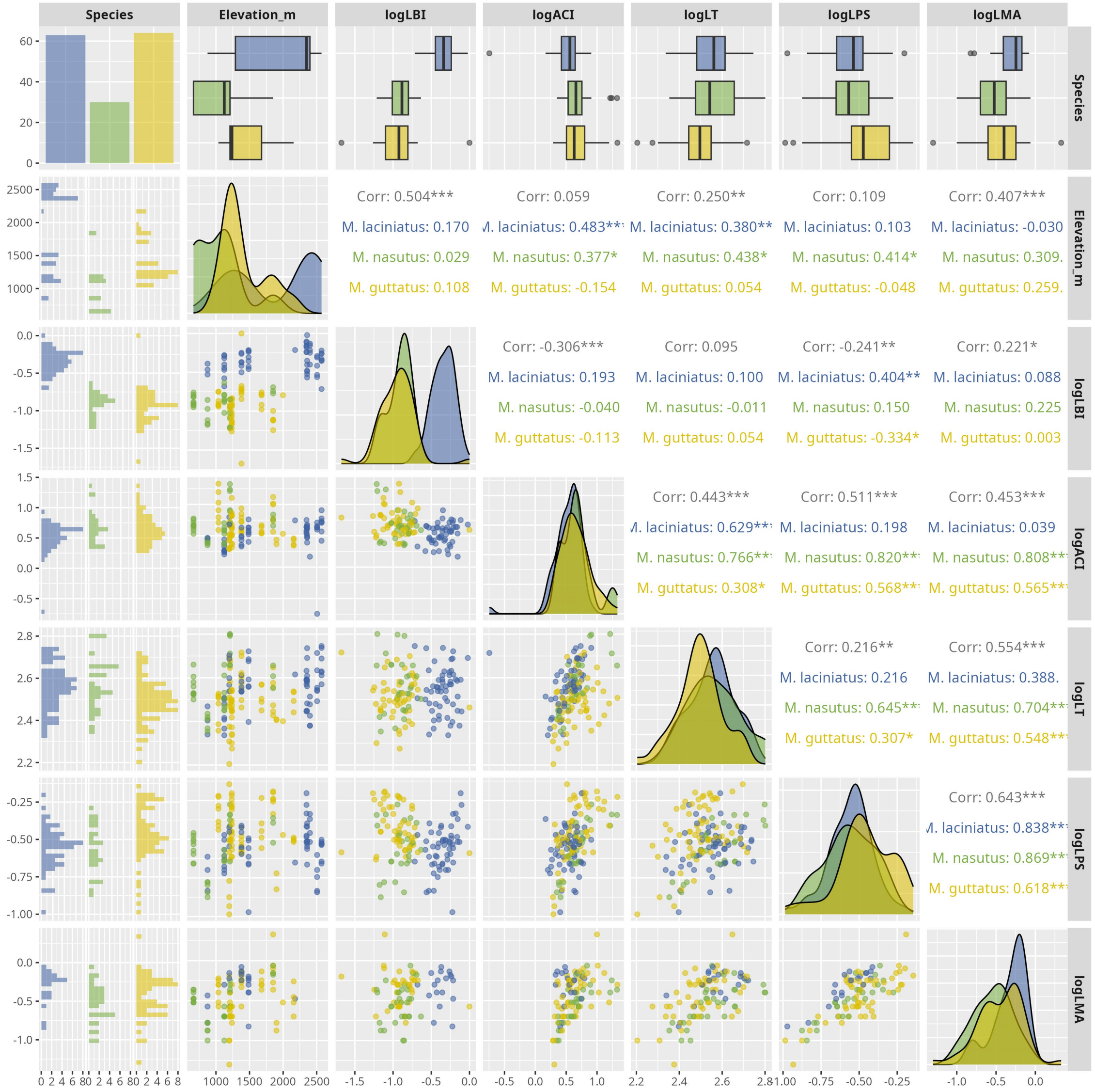


**Figure S1**. Correlation matrix of log-transformed leaf functional traits by host species. The plot is based on the Spearman's rho. Significance levels are represented by *ns* (not significant) and asterisks [$p$ < .05 (*), $p$ < 0.01 (**), $p$ < .001 (***), and $p$ < .0001 (****)].

### 13.2 Figure S2

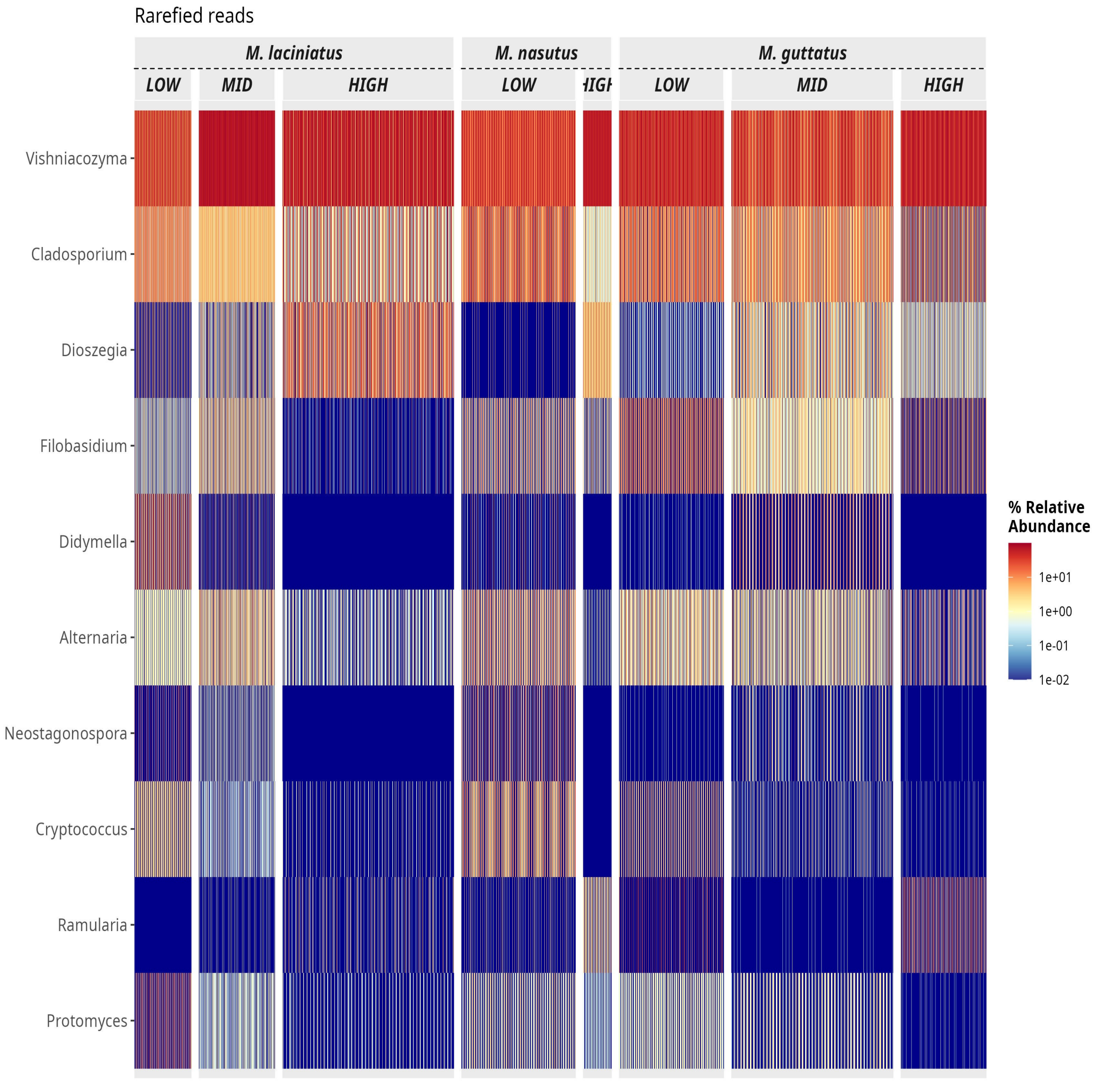


**Figure S2**. Heatmap of the top 10 most abundant FEF genera in the dataset. The heatmap is faceted by host species and elevation zone.

#### 13.3 Figure S3

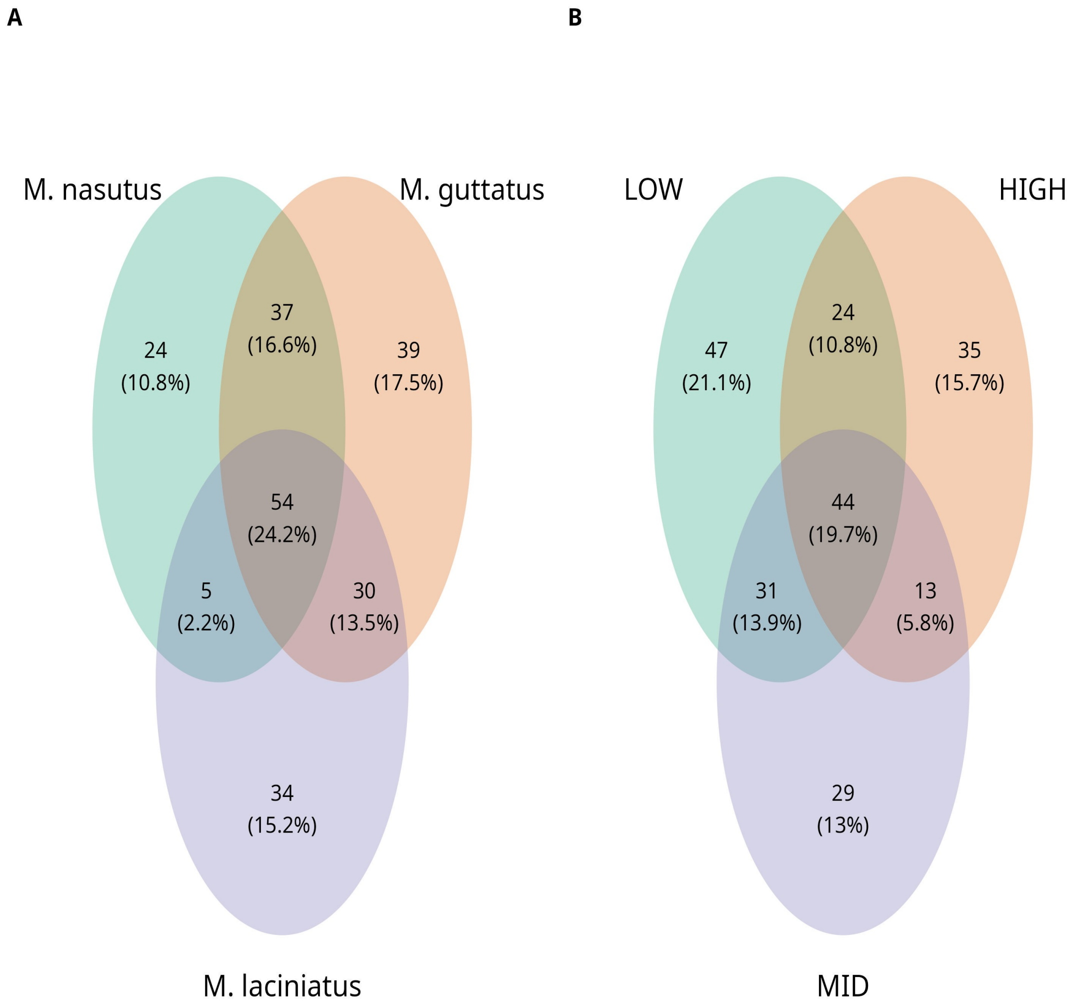


**Figure S3**. Venn diagrams of presence and absence of ASVs among species and elevation zones un-rarified data and rarified data. A) Represents the overlap of ASVs in in host species and B) per elevation zone. Number of ASVs are represented the count and in parentheses as a ratio shared between host species and elevation zone

#### 13.4 Figure S4

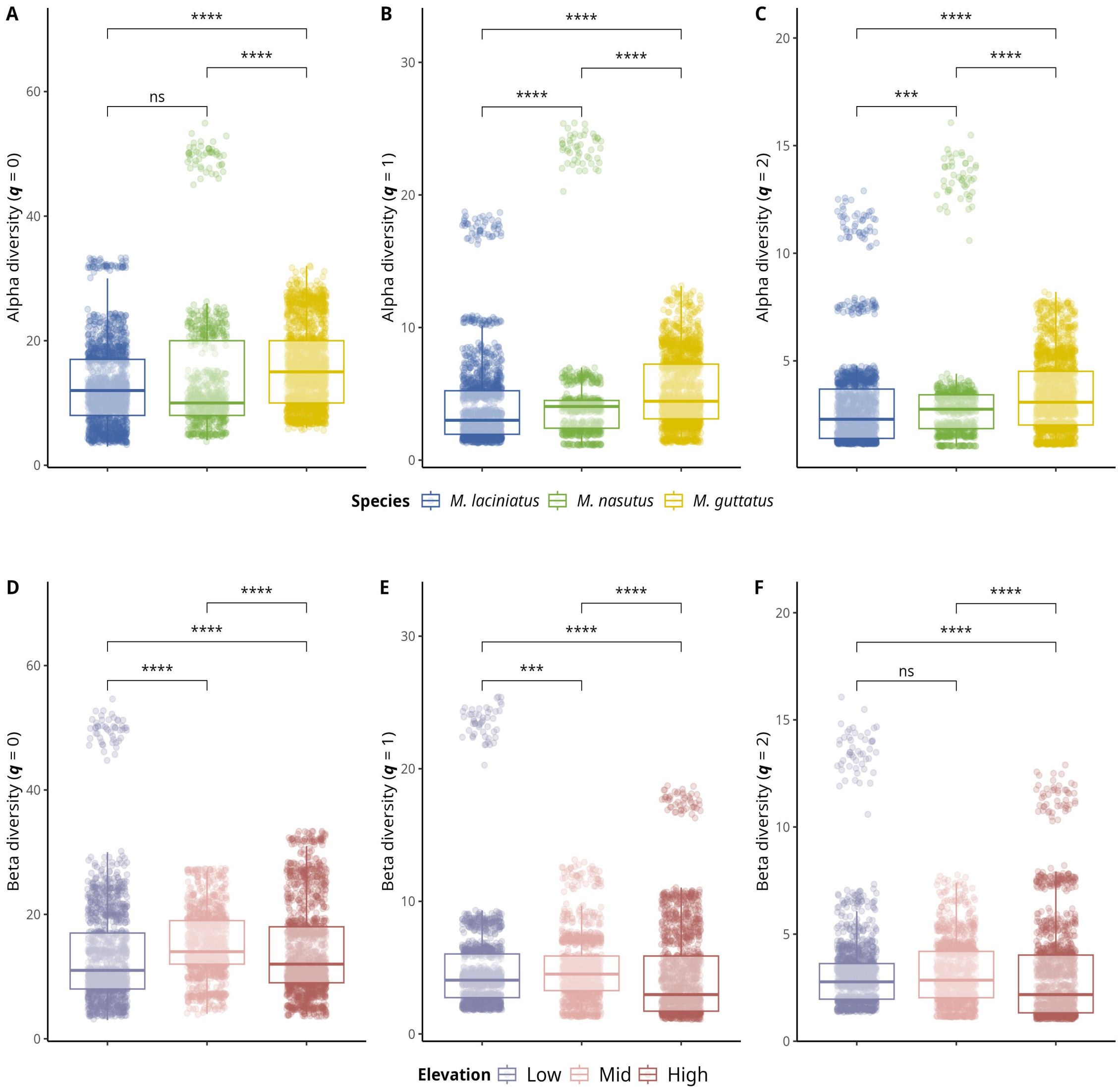


**Figure S4**. Alpha and beta diversity means comparisons in host species and elevation zones. Alpha diversity means comparisons; A) Observed ASV richness ($q$ = 0); B) Shannon's entropy ($q$ = 1); and C) Inverse Simpson's index ($q$ = 2) per host species. A-C) Blue filled boxplots correspond to *M. laciniatus*, green filled to *M. nasutus*, and yellow to *M. guttatus*. Beta diversity mean comparisons; A) Observed ASV richness ($q$ = 0); B) Shannon's entropy ($q$ = 1); and C) Inverse Simpson's index ($q$ = 2) per elevation zone. D-F) Violet boxplots correspond to LOW elevation sites, pink filled to MID and light maroon to HIGH elevation sites, while squares represent *M. laciniatus*, circles *M. nasutus* and triangles *M. guttatus*. Significance levels are represented by *ns* (not significant) and asterisks [$p$ < .05 (*), $p$ < .01 (**), $p$ < .001 (***), and $p$ < .0001 (****)].

### 13.5 Figure S5

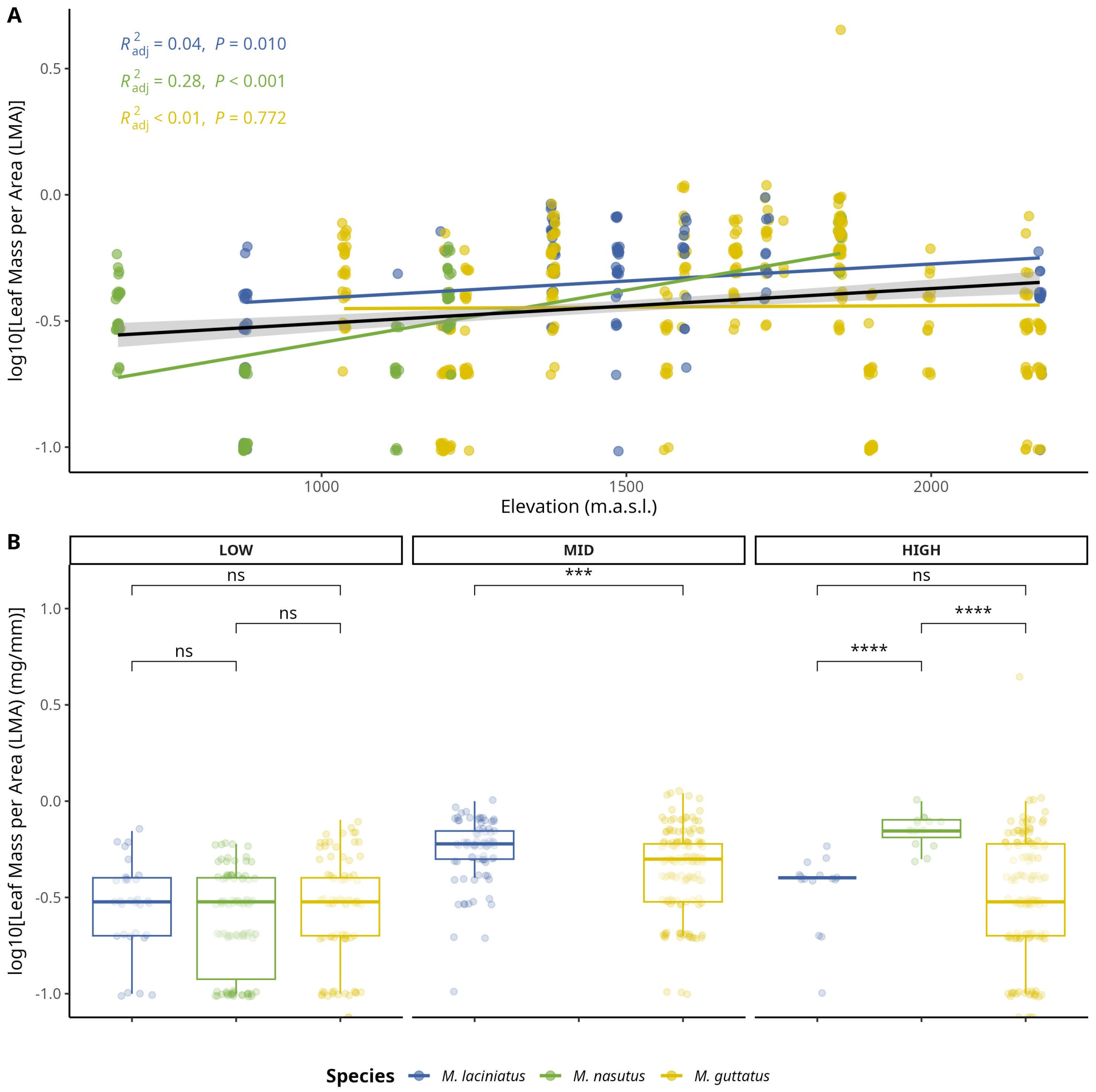


**Figure S5**. Comparison of log-transformed leaf mass per area (LMA) by species and elevation zone. A) Compares logLMA means by species and elevation zone: LOW (<1220 m), MID (1221 - 1828 m), HIGH (> 1829 m). B) Change in logLMA per species as elevation increases. The black trend line represents the fitted model for all data points, see main text for $R^2_{adj}$ and $p$ values. Significance levels are represented by *ns* (not significant) and asterisks [$p < .05$ (*), $p < 0.01$ (**), $p < .001$ (***), and $p < .0001$ (****)].

#### 13.6 Figure S6

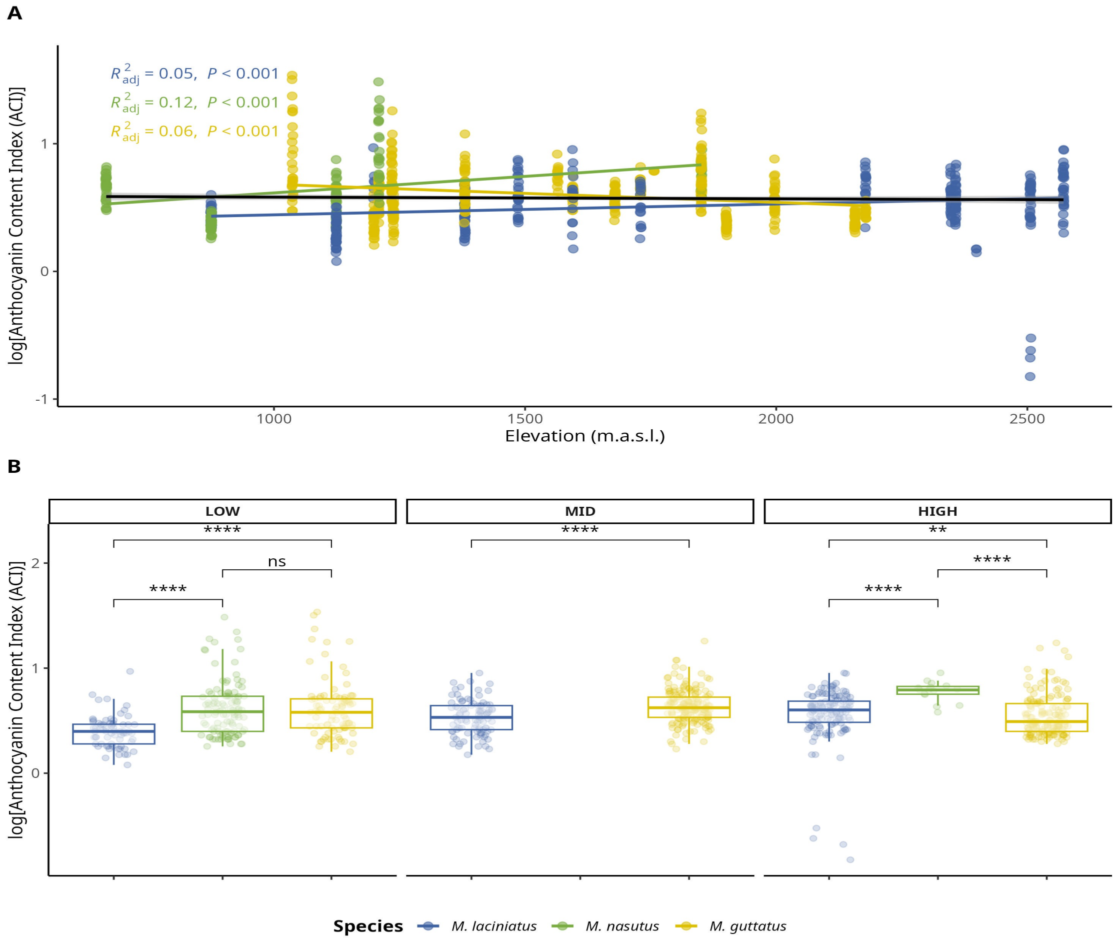


**Figure S6**. Change in log-transformed anthocyanin content index (ACI) by species and elevation zone. A) Compares logACI means by species and elevation zone: LOW (<1220 m), MID (1221 - 1828 m), HIGH (> 1829 m). The black trend line represents the fitted model for all data points, see main text for $R^2_{adj}$ and $p$ values. B) Change in logLPS per species as elevation increases. Significance levels are represented by *ns* (not significant) and asterisks [$p < .05$ (*), $p < .01$ (**), $p < .001$ (***), and $p < .0001$ (****)].

#### 13.7 Figure S7

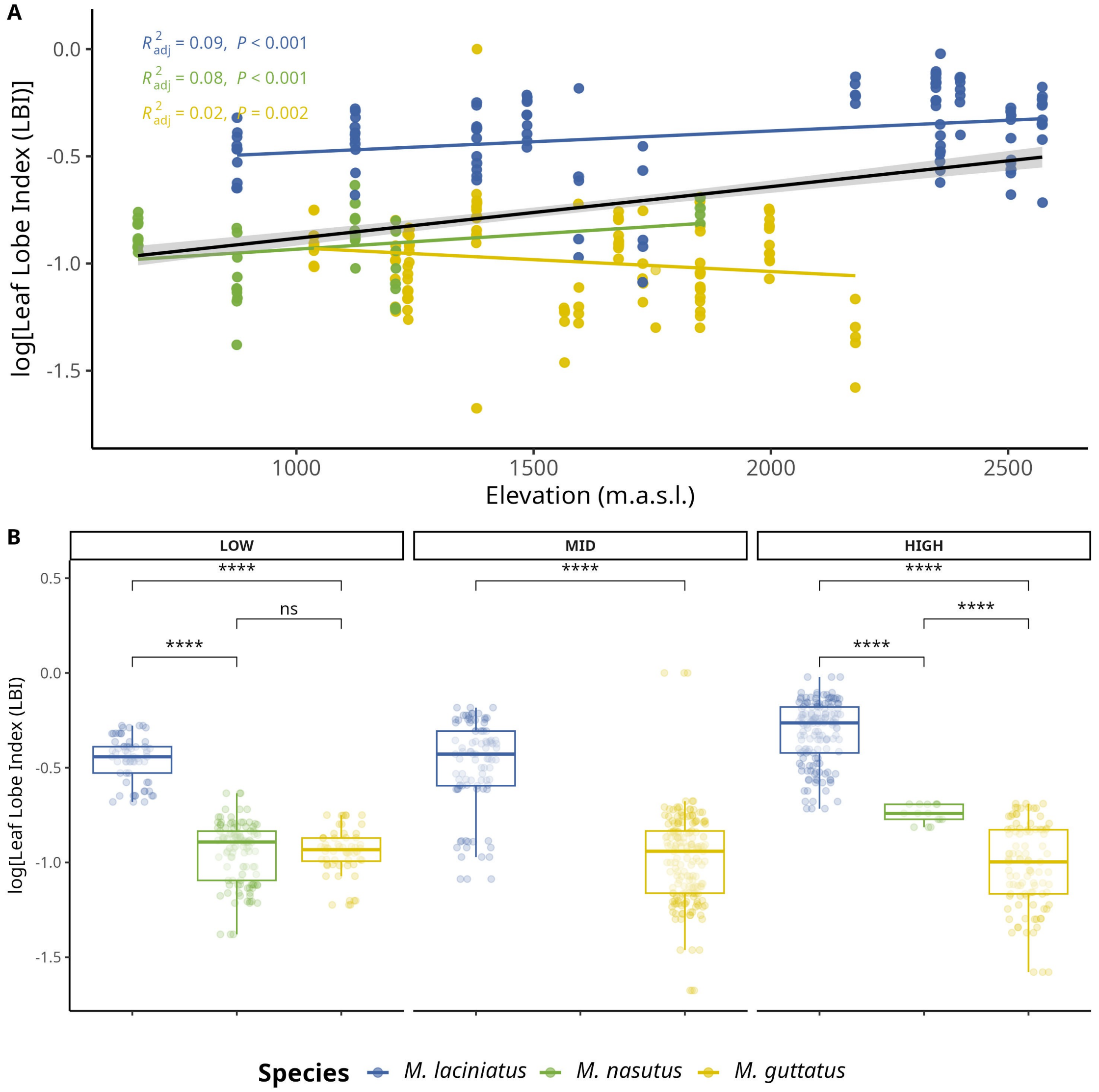


**Figure S7**. Change in log-transformed leaf lobe index (LBI) by species and elevation zone. A) Compares logLBI means by species and elevation zone: LOW (<1220 m), MID (1221 - 1828 m), HIGH (> 1829 m). The black trend line represents the fitted model for all data points, see main text for $R^2_{adj}$ and *p* values. B) Change in logLPS per species as elevation increases. Significance levels are represented by *ns* (not significant) and asterisks [*p* < .05 (*), *p* < .01 (**), *p* < .001 (***), and *p* < .0001 (****)].

#### 13.8 Figure S8

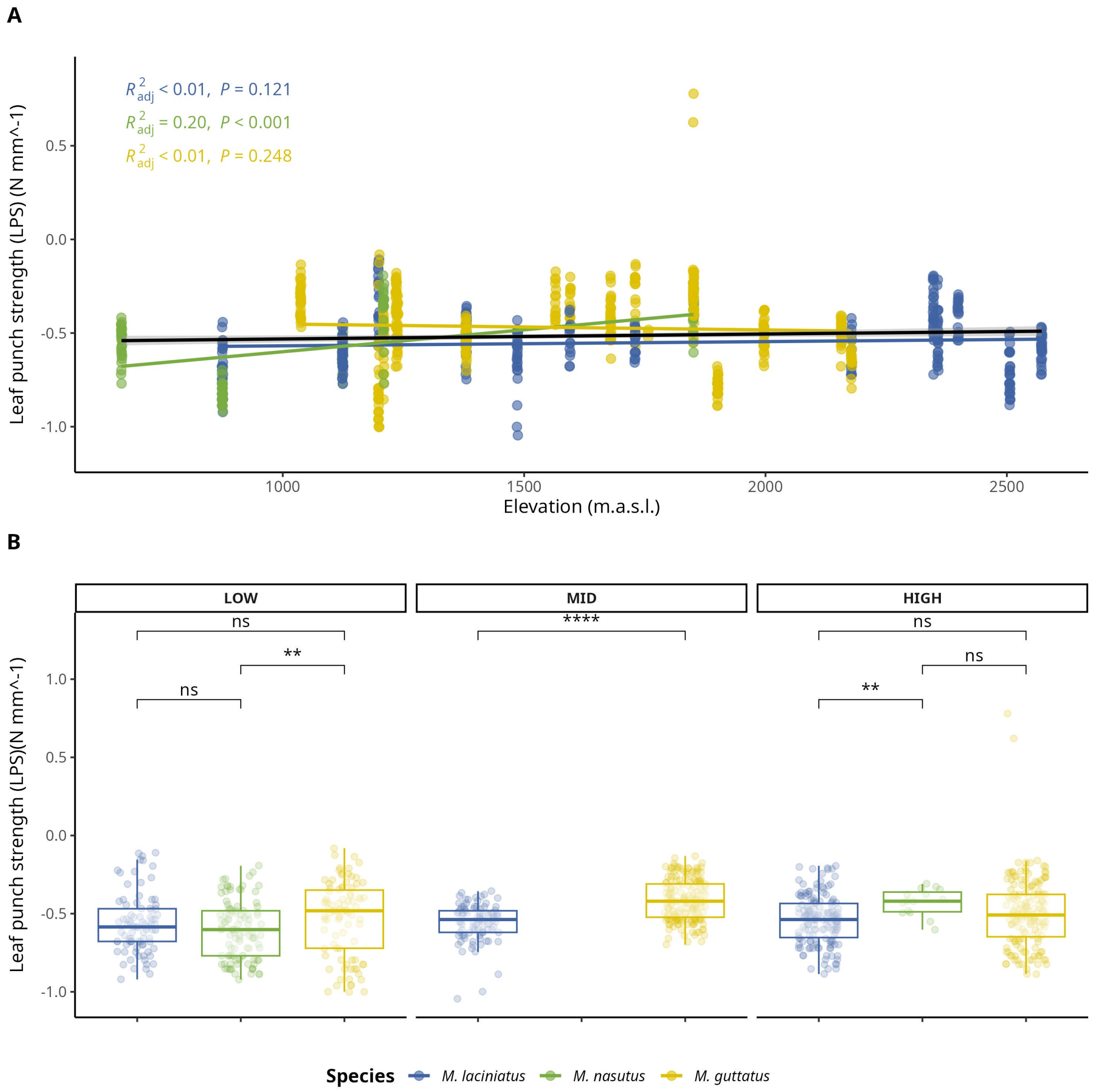


**Figure S8**. Change in log-transformed leaf-punch strength (LPS), a measure of leaf toughness, by species and elevation zone. A) Compares logLPS means by species and elevation zone: LOW (<1220 m), MID (1221 - 1828 m), HIGH (> 1829 m). The black trend line represents the fitted model for all data points, see main text for $R^2_{adj}$ and $p$ values. B) Change in logLPS per species as elevation increases. Significance levels are represented by *ns* (not significant) and asterisks [$p < .05$ (*), $p < .01$ (**), $p < .001$ (***), and $p < .0001$ (****)].

### 13.9 Figure S9

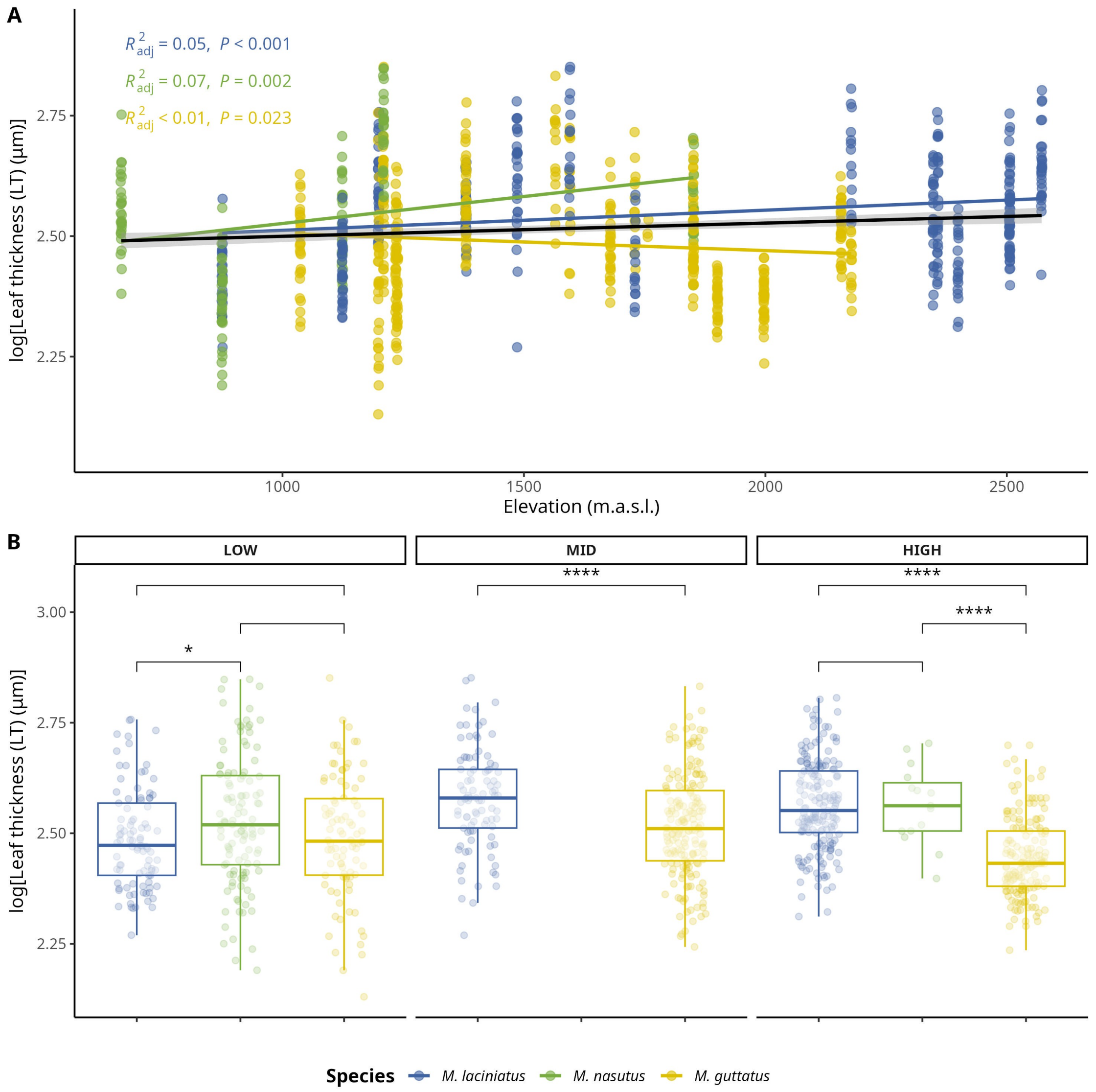


**Figure S9**. Change in log-transformed leaf thickness (LT) ( m) by species and elevation zone. A) Compares logLT means by species and elevation zone: LOW (<1220 m), MID (1221 - 1828 m), HIGH (> 1829 m). The black trend line represents the fitted model for all data points, see main text for $R^2_{adj}$ and *p* values. B) Change in logLT per species as elevation increases. Significance levels are represented by *ns* (not significant) and asterisks [*p* < .05 (*), *p* < .01 (**), *p* < .001 (***), and *p* < .0001 (****)].

#### 13.10 Table S1

**Number of individuals collected per population/site and species.**

| Site | Site Name | Forest/Park | Latitude | Longitude | *M. laciniatus* | *M. nasutus* | *M. guttatus* |
|---|---|---|---|---|---|---|---|
| AGFB | Angel Falls | Sierra National Forest | 37.338230 | -119.570677 | 11 | 10 | |
| BHEB | Bald Mountain | Sierra National Forest | 37.053544 | -119.393233 | | 9 | 10 |
| BMLZ | Bretz Mill Road | Sierra National Forest | 37.090625 | -119.314015 | 5 | | 5 |
| CCNB | Crab Tree Road | Stanislaus National Forest | 38.183927 | -119.963252 | | | 12 |
| CCRB | Carson Creek | Mariposa County | 37.486337 | -119.997920 | | 9 | |
| CFZZ | Crane Flat | Yosemite National Park | 37.755986 | -119.802952 | | | 10 |
| HDRB | Highland Drive Road | Sierra National Forest | 37.288671 | -119.534709 | | | 10 |
| HHBB | Hetch Hetchy B | Yosemite National Park | 37.959056 | -119.778553 | 10 | | 10 |
| HULB | Huntingto n Lake | Sierra National Forest | 37.235530 | -119.149620 | 5 | | 5 |
| HUOB | Huntingto n Lake Overview | Sierra National Forest | 37.150723 | -119.273507 | | 5 | 17 |
| KNOB | The Knobs | Yosemite National Park | 37.853852 | -119.440369 | 10 | | |
| LMZZ | Little Meadow | Yosemite National Park | 37.767781 | -119.772719 | | | 10 |

| | | | | | | | |
|---|---|---|---|---|---|---|---|
| NFWB | North Fork Willow Creek | Sierra National Forest | 37.257397 | -119.521415 | 10 | 10 | |
| OPNZ | Olmstead Point | Yosemite National Park | 37.807232 | -119.486313 | 20 | | |
| OSKB | Onion Skins | Yosemite National Park | 37.842693 | -119.594572 | 7 | | |
| SDSB | Stoddard Spring | Stanislaus National Forest | 38.136194 | -120.094614 | | | 10 |
| SHLZ | Shaver Lake | Sierra National Forest | 37.145131 | -119.307426 | 5 | | 5 |
| SPLB | Spotless | Yosemite National Park | 37.847194 | -119.586129 | 9 | | |
| TRTZ | Turtle Back Dome | Yosemite National Park | 37.716098 | -119.705317 | 10 | | |
| TWGB | Tawonga | Stanislaus National Forest | 37.858820 | -119.951305 | | | 9 |
| WCFB | Wildcat Falls | Yosemite National Park | 37.723593 | -119.719077 | 11 | | 12 |
| WSHB | Wawona School | Yosemite National Park | 37.543343 | -119.647878 | | | 12 |
| YCNB | Yosemite Creek B | Yosemite National Park | 37.843904 | -119.572754 | 10 | | |

#### 13.11 Table S2

***Prevalence of phyla in samples per species.***

| | | Occurrence |
|---|---|---|
| *M. laciniatus* | | |
| | Ascomycota | 57 |
| | Basidiomycota | 62 |
| | Mortierellomycota | 2 |
| | Olpidiomycota | 2 |
| *M. nasutus* | | |
| | Ascomycota | 29 |
| | Basidiomycota | 27 |
| | Chytridiomycota | 2 |
| | Mortierellomycota | 2 |
| | Rozellomycota | 2 |
| *M. guttatus* | | |
| | Ascomycota | 58 |
| | Basidiomycota | 63 |
| | Mortierellomycota | 2 |
| | Olpidiomycota | 1 |

Occurrence represents the number of samples where each phylum was detected per species.

#### 13.12 Table S3

| Term | Df | Sum. of Sqs. | F-value | p-value |
|---|---|---|---|---|
| logLBI | 1.000 | 57.693 | 306.561 | *** |
| Elevation_m | 1.000 | 53.745 | 285.579 | *** |
| Species | 2.000 | 29.173 | 77.508 | *** |
| Elevation_m:Species | 2.000 | 45.614 | 121.188 | *** |
| *Residual* | 3,443.000 | 647.959 | | |

Significance levels are represented by ns (not significant) and asterisks [p < .05 (*), p < .01 (**), p < .001 (***), and p < .0001 (****)].

#### 13.13 Supplementary methods

##### 13.13.1 Illumina TruSeq adapters

The modified primers for the first PCR (adapter ligation and ITS1 amplification) were as follows:

5’-CAC TCT TTC CCT ACA CGA CGC TCT TCC GAT CTC TTG GTC ATT TAG AGG AAG

TAA-3’ (forward) and 5’-GTG ACT GGA GTT CAG ACG TGT GCT CTT CCG ATC TGC TGC GTT CTT CAT CGA TGC-3’ (reverse). See (Gardes and Bruns, 1993) and (White, T. J. et al., 1990) for more details.
